# Na_V_1.7-dependent peripheral sensitization drives chronic pain in Parkinson’s disease

**DOI:** 10.64898/2026.09.09.750469

**Authors:** Tyler S. Nelson, Naomi K. Grabus, Heather N. Allen, Aida Calderon-Rivera, Santiago Loya-Lopez, Erick J. Rodriguez-Palma, Alaina L. Waters, Eslam Elhanafy, Stephanie I. Shiers, Ishwarya Sankaranarayanan, Kimberly Gomez, Theodore J. Price, Rajesh Khanna

## Abstract

Pain is among the most prevalent and disabling nonmotor symptoms of Parkinson’s disease (PD), yet its mechanisms remain poorly defined and effective treatments are limited. Safinamide is one of the few drugs reported to improve pain in PD, but the mechanism underlying this effect is unknown. Here, we show that nigrostriatal neurodegeneration produces persistent hyperexcitability of primary sensory neurons associated with dysregulation of the voltage-gated sodium channel Na_V_1.7. In a brain-restricted 6-hydroxydopamine (6-OHDA) model, small-diameter dorsal root ganglion (DRG) neurons exhibited increased sodium current density and altered voltage-dependent inactivation, with the excess current eliminated by selective Na_V_1.7 blockade. Safinamide directly inhibited a Na_V_1.7-dependent component of sensory neuron sodium current and reversed established pain-like behaviors. Pharmacological disruption of Na_V_1.7 regulation by collapsin response mediator protein 2 (CRMP2) normalized DRG hyperexcitability and reversed mechanical and thermal hypersensitivity, whereas genetic disruption of the Na_V_1.7 CRMP2 regulatory sequence prevented the development of 6-OHDA-induced pain-like behaviors for up to 30 weeks despite preservation of the Parkinsonian motor phenotype. Transcriptomic profiling of human PD DRGs revealed limited global transcriptional remodeling with selective alterations in genes associated with sensory neuron excitability. Together, these findings demonstrate that dopaminergic neurodegeneration initiated within the brain is sufficient to drive persistent peripheral sensory neuron dysfunction and identify CRMP2-dependent regulation of Na_V_1.7 as a therapeutic target for Parkinsonian pain.

## Introduction

Parkinson’s disease (PD) is the second most common neurodegenerative disorder, affecting more than 10 million individuals worldwide (1, 2). Although PD is classically defined by progressive degeneration of nigrostriatal dopaminergic neurons and disabling motor dysfunction, non-motor symptoms are increasingly recognized as major determinants of patient quality of life (3–5). Chronic pain affects up to 85% of individuals with PD, can precede the onset of motor symptoms, and is associated with impaired mobility, depression, sleep disturbance, and reduced quality of life (6–10). Despite its prevalence and clinical burden, the mechanisms underlying PD-associated pain remain poorly understood, and effective treatments remain limited.

The pathophysiology of PD pain has traditionally been attributed to altered central nociceptive processing resulting from degeneration of dopaminergic pathways (11, 12). Although central mechanisms undoubtedly contribute, dopamine loss alone does not fully explain the clinical features of Parkinsonian pain. Pain severity correlates incompletely with motor dysfunction, responses to dopaminergic replacement therapy are variable, and pain frequently persists despite optimization of motor symptoms (7, 13). Notably, safinamide (Xadago®), a monoamine oxidase-B (MAO-B) inhibitor approved as an adjunct therapy for PD, is among the few pharmacological treatments consistently reported to improve pain-related outcomes in patients with PD (14–18). However, despite its demonstrated clinical efficacy, the mechanism underlying its analgesic effects remains unknown. In addition to MAO-B inhibition, safinamide modulates voltage-gated sodium and calcium channels (19, 20), raising the possibility that its analgesic effects involve mechanisms outside canonical dopaminergic signaling.

Growing evidence indicates that PD pathology extends beyond the central nervous system (21). Peripheral abnormalities, including autonomic dysfunction, enteric pathology, small fiber neuropathy, and α-synuclein accumulation in peripheral neurons, can occur during the course of disease and may contribute to non-motor symptoms (22–25). More recently, Zhang et al. demonstrated increased excitability and sodium current density in small diameter dorsal root ganglion (DRG) neurons following systemic methyl-4-phenyl-1,2,3,6-tetrahydropyridine (MPTP) administration, accompanied by increased expression of *Scn9a* and *Scn10a* and suppression of DRG hyperexcitability by safinamide (26). These findings provided important evidence that primary sensory neuron dysfunction can accompany experimental Parkinsonism. However, because MPTP was administered systemically, it remains unclear whether the peripheral phenotype arose downstream of nigrostriatal neurodegeneration or from direct or indirect peripheral consequences of toxin exposure. The sodium channel(s) responsible for the excess current and the ensuing peripheral hyperexcitability remain unknown.

The voltage-gated sodium channel Na_V_1.7 is highly enriched in nociceptive sensory neurons and is a major determinant of action potential initiation and pain signaling (27). Loss of function variants in *SCN9A* cause congenital insensitivity to pain, whereas gain-of-function variants produce severe inherited pain syndromes, establishing a direct relationship between Na_V_1.7 function and human pain perception (28–32). Importantly, two independent studies reported *SCN9A* variants with increased pain susceptibility in PD patients, providing human genetic support for a potential contribution of Na_V_1.7 to Parkinsonian pain (33, 34). Na_V_1.7 function is also regulated by collapsin response mediator protein 2 (CRMP2), which controls channel trafficking and membrane availability through a discrete CRMP2 regulatory sequence (CRS) within Na_V_1.7, and disruption of this interaction reduces Na_V_1.7 current and pathological pain (35–38). However, whether Na_V_1.7 function is altered in peripheral sensory neurons following nigrostriatal neurodegeneration, whether CRMP2-dependent regulation contributes to this process, and whether these peripheral mechanisms are required for PD-associated pain remain unknown.

Here, we used a brain-restricted 6-hydroxydopamine (6-OHDA) lesion to test if nigrostriatal neurodegeneration is sufficient to induce persistent peripheral sensory neuron sensitization and whether Na_V_1.7 contributes to this process. We demonstrate persistent hyperexcitability of primary sensory neurons and identify increased Na_V_1.7-dependent current as a major component of this phenotype. We show that safinamide directly inhibits a Na_V_1.7-dependent component of sensory neuron sodium current and reverses pain-like behaviors following nigrostriatal degeneration. Pharmacological disruption of CRMP2-dependent Na_V_1.7 regulation normalized DRG neuron hyperexcitability and reversed established pain, whereas genetic disruption of the Na_V_1.7 CRS prevented development of 6-OHDA-induced hypersensitivity for up to 30 weeks. Finally, transcriptomic analysis of human PD DRG revealed limited global transcriptional remodeling, with unchanged *SCN9A* and *DPYSL2* expression but altered expression patterns across HCN channel family members. Together, these findings establish that nigrostriatal neurodegeneration can drive persistent peripheral sensory neuron dysfunction and identify CRMP2-dependent regulation of Na_V_1.7 as a potential therapeutic target for Parkinsonian pain.

## Results

### Safinamide inhibits a NaV1.7-dependent component of sodium current in primary sensory neurons

Safinamide directly modulates voltage-gated sodium channels independently of MAO-B inhibition (19, 20), and suppresses hyperexcitability in DRG neurons from MPTP-treated mice (26), but the responsible channel subtype is unknown. To determine whether Na_V_1.7 contributes to the actions of safinamide on sensory neuron sodium currents, we performed whole-cell voltage-clamp recordings from small-diameter DRG neurons isolated from wild-type mice under four bath conditions: vehicle, safinamide (10 μM), the selective Na_V_1.7 inhibitor Protoxin-II (ProTx-II; Tocris/Bio-Techne, Cat#4023, 5 nM) (39), or safinamide and ProTx-II (**Fig. 1A**). Representative sodium current traces are shown in **Figure 1B**. Analysis of current-voltage relationships demonstrated that both safinamide and ProTx-II reduced sodium current density relative to vehicle (**Fig. 1C**). Peak sodium current density was significantly reduced by safinamide (∼51% inhibition) and ProTx-II (∼37% inhibition), whereas combined treatment produced ∼58% inhibition, which was not significantly greater than that produced by ProTx-II alone (**Fig. 1D**). These findings indicate that safinamide inhibits a ProTx-II-sensitive, Na_V_1.7-dependent component of the total sodium current in primary sensory neurons.

**Figure 1.**
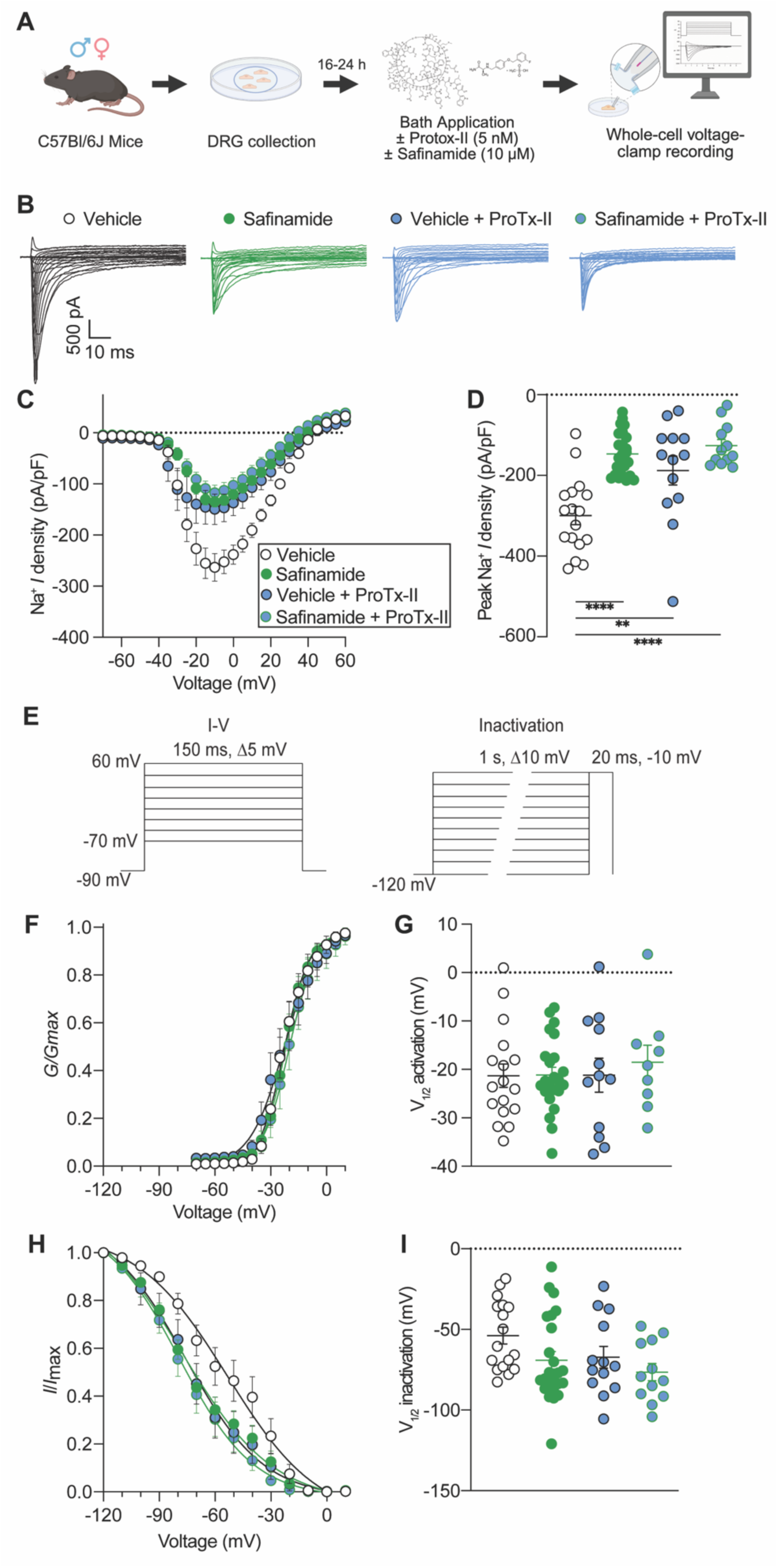
Safinamide inhibits a ProTx-II-sensitive component of voltage-gated sodium current in primary sensory neurons. **(A)** Experimental design for whole-cell voltage-clamp recordings from dorsal root ganglion (DRG) neurons isolated from naïve C57BL/6J mice. DRG neurons were cultured for 16-24 h and treated during recording with vehicle (0.1% DMSO), safinamide (10 μM), ProTx-II (5 nM), or safinamide + ProTx-II. **(B)** Representative whole-cell sodium current traces from each treatment condition. **(C)** Current–voltage (I–V) relationships for total voltage-gated sodium current following vehicle, safinamide, ProTx-II, or safinamide + ProTx-II treatment. **(D)** Peak sodium current density for each treatment condition. Safinamide and ProTx-II significantly reduced peak sodium current density. **(E)** Voltage-clamp protocols used to determine sodium current activation and steady-state inactivation. **(F)** Voltage dependence of activation for each treatment condition. **(G)** Half-maximal voltage of activation (V_1/2_ activation) derived from Boltzmann fits. **(H)** Voltage dependence of steady-state inactivation for each treatment condition. **(I)** Half-maximal voltage of inactivation (V_1/2_ inactivation) derived from Boltzmann fits. For **D, G,** and **I**, statistical comparisons were performed using two-way ANOVA followed by Tukey’s multiple comparisons test. N = 9-24 cells/group. Data are presented as mean ± SEM with individual cells shown where applicable. **P < 0.01, ****P < 0.0001.

We next examined whether safinamide altered the voltage dependence of sodium channel gating using activation and steady-state inactivation protocols (**Fig. 1E**). Safinamide, ProTx-II, and their combination produced only modest effects on the voltage dependence of activation (**Fig. 1F**), with no significant differences in V_1/2_ activation (**Fig. 1G**). Steady-state inactivation was likewise not significantly altered by treatment (**Fig. 1H**), with no significant differences in V_1/2_ inactivation (**Fig. 1I**). Together, these electrophysiological findings support inhibition of a Na_V_1.7-dependent component of sensory neuron sodium current by safinamide without substantial alteration of voltage-dependent gating. Molecular docking was consistent with this interpretation: across five independent NaV1.7 cryo-EM structures, the top-ranked safinamide poses consistently occupied the central pore cavity and engaged the DIV-S6 residue F1748 through π-stacking (**Supplementary Fig. S1** and **Supplementary Table S1**), the phenylalanine that anchors the canonical local anesthetic receptor site. Convergence of the pharmacology, a safinamide-sensitive current that is also ProTx-II-sensitive and shows no gating shift, together with a conserved pore-blocking pose predicted in silico identifies Na_V_1.7 as the candidate mediator of safinamide-sensitive sodium current.

### Unilateral 6-OHDA lesions produce nigrostriatal dopaminergic degeneration and motor dysfunction

To model Parkinsonian neurodegeneration, mice received unilateral injection of 6-OHDA into the right medial forebrain bundle (MFB), a well-established model of nigrostriatal dopaminergic degeneration (**Fig. 2A**) (40, 41). Three weeks after 6-OHDA administration, immunohistochemical analysis demonstrated a marked loss of tyrosine hydroxylase (TH) immunoreactivity in the ipsilateral striatum compared with the contralateral hemisphere and sham controls, confirming robust disruption of nigrostriatal dopaminergic innervation (**Fig. 2, B and C**). We next assessed the functional consequences of the lesion using apomorphine-induced rotational behavior (**Fig. 2D**). 6-OHDA-lesioned mice exhibited robust contralateral rotations, consistent with substantial unilateral dopaminergic denervation (**Fig. 2E**). Motor function was further evaluated using the accelerating rotarod, where 6-OHDA mice exhibited reduced latency to fall and lower rotational speed at the time of fall compared with sham controls (**Fig. 2, F and G**). Finally, 6-OHDA mice exhibited impaired nest-building behavior (**Fig. 2, H and I**). Together, these histological and behavioral measures confirm a robust unilateral Parkinsonian lesion characterized by nigrostriatal dopaminergic degeneration and persistent motor dysfunction.

**Figure 2.**
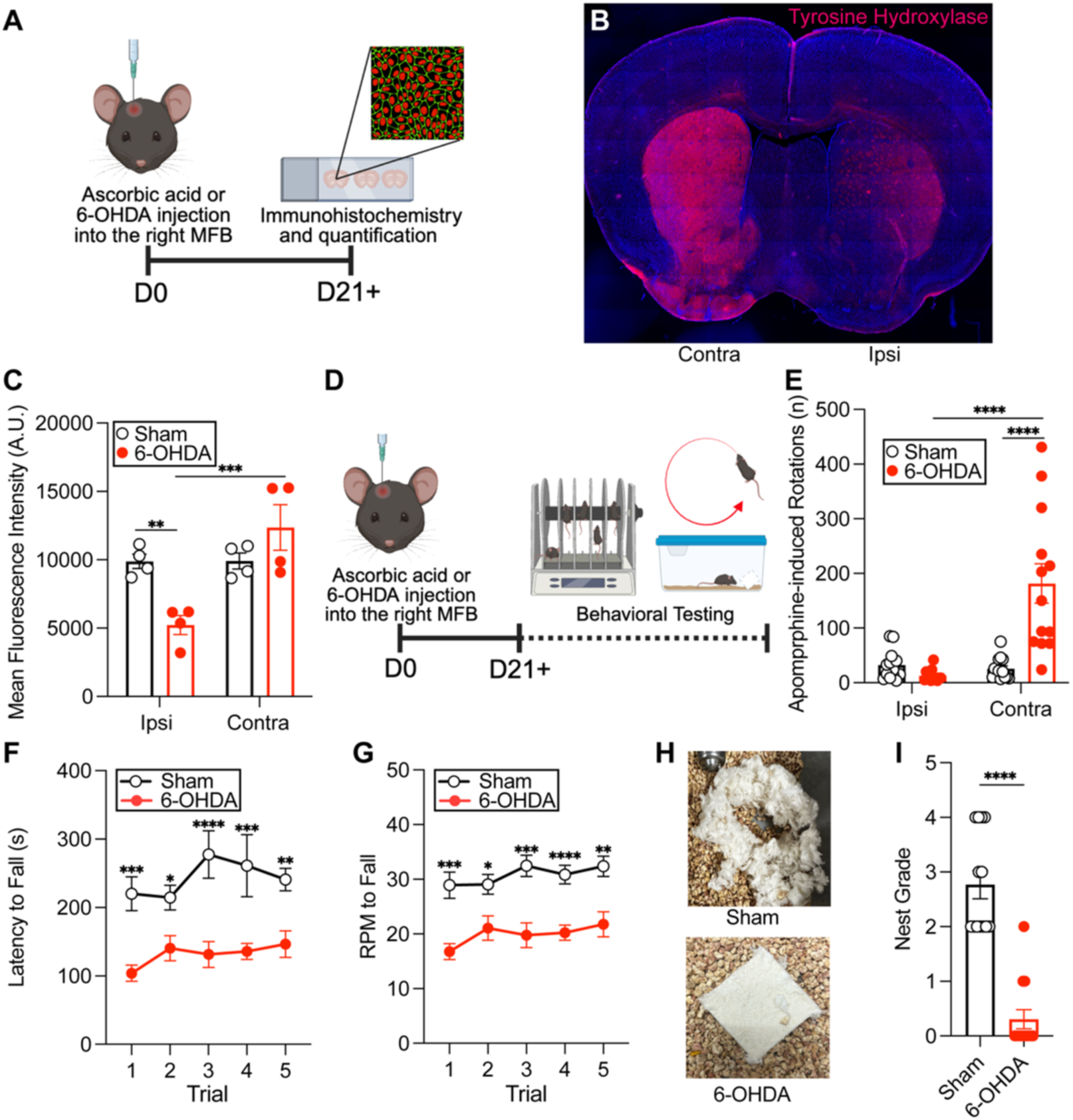
Unilateral 6-OHDA lesions produce nigrostriatal dopaminergic degeneration and motor dysfunction. **(A)** Experimental timeline for unilateral injection of ascorbic acid (sham) or 6-hydroxydopamine (6-OHDA) into the right medial forebrain bundle (MFB), followed by tyrosine hydroxylase (TH) immunohistochemistry and quantification at ≥21 days post-lesion. **(B)** Representative coronal brain section showing TH immunoreactivity (magenta) following unilateral 6-OHDA lesion, demonstrating loss of dopaminergic innervation in the ipsilateral striatum relative to the contralateral hemisphere. **(C)** Quantification of mean TH fluorescence intensity in the ipsilateral (Ipsi) and contralateral (Contra) striatum of sham and 6-OHDA mice (n = 4 mice/group). Statistical comparisons were performed using two-way ANOVA followed by Tukey’s multiple comparisons test. **(D)** Experimental timeline for behavioral characterization beginning ≥21 days following unilateral MFB injection. **(E)** Apomorphine-induced rotational behavior in sham and 6-OHDA mice (n = 13 mice/group), demonstrating robust contralateral rotations following 6-OHDA lesion. Statistical comparisons were performed using two-way ANOVA followed by Uncorrected Fisher’s LSD. **(F)** Accelerating rotarod latency to fall and **(G)** rotational speed (RPM) at the time of fall across five trials in sham and 6-OHDA mice (n = 13 mice/group). For **F–G**, statistical comparisons were performed using repeated-measures two-way ANOVA followed by Tuk multiple comparisons test. **(H)** Representative nests constructed by sham (above) and 6-OHDA mice (below). **(I)** Quantification of nest-building performance in sham and 6-OHDA mice (n = 13 mice/group). Statistical comparison was performed using an unpaired two-tailed Student’s *t* test. Data are presented as mean ± SEM. * P < 0.05, ** P < 0.01, *** P < 0.001, **** P < 0.0001.

### Safinamide rapidly reverses established pain-like behaviors in experimental Parkinson’s disease

Having validated the lesion model, we next asked whether unilateral 6-OHDA lesions produced pain-like behaviors and, if so, whether safinamide reversed them. Previous studies have demonstrated that bilateral sensory hypersensitivity emerges within the first week following unilateral 6-OHDA lesion and remains persistently elevated thereafter (40, 42). Consistent with this established time course, 6-OHDA-treated mice developed significant mechanical hypersensitivity, dynamic mechanical allodynia, and cold hypersensitivity by 10 days following lesioning, whereas sham-operated mice exhibited no changes in sensory behavior (**Fig. 3, A-D**). We therefore used ≥10 days post-lesion as the experimental window for subsequent studies examining Parkinsonian pain. To determine the analgesic efficacy of safinamide, mice received a single intraperitoneal injection of vehicle or safinamide (100 mg/kg) (26), and sensory behaviors were assessed over the subsequent two hours (**Fig. 3E**). Safinamide rapidly and significantly reversed mechanical hypersensitivity in 6-OHDA mice, restoring withdrawal thresholds to sham levels within 60 minutes before the effect diminished by 120 minutes (**Fig. 3F**). Similarly, safinamide significantly reduced brush-evoked responses and acetone-evoked withdrawal duration at the 60-minute time point, indicating reversal of dynamic mechanical allodynia and cold hypersensitivity, respectively (**Fig. 3, G-H**). Safinamide did not alter sensory responses in sham-operated mice. Together, these findings show that acute safinamide administration effectively reverses multiple modalities of established pain-like behavior in experimental PD.

**Figure 3.**
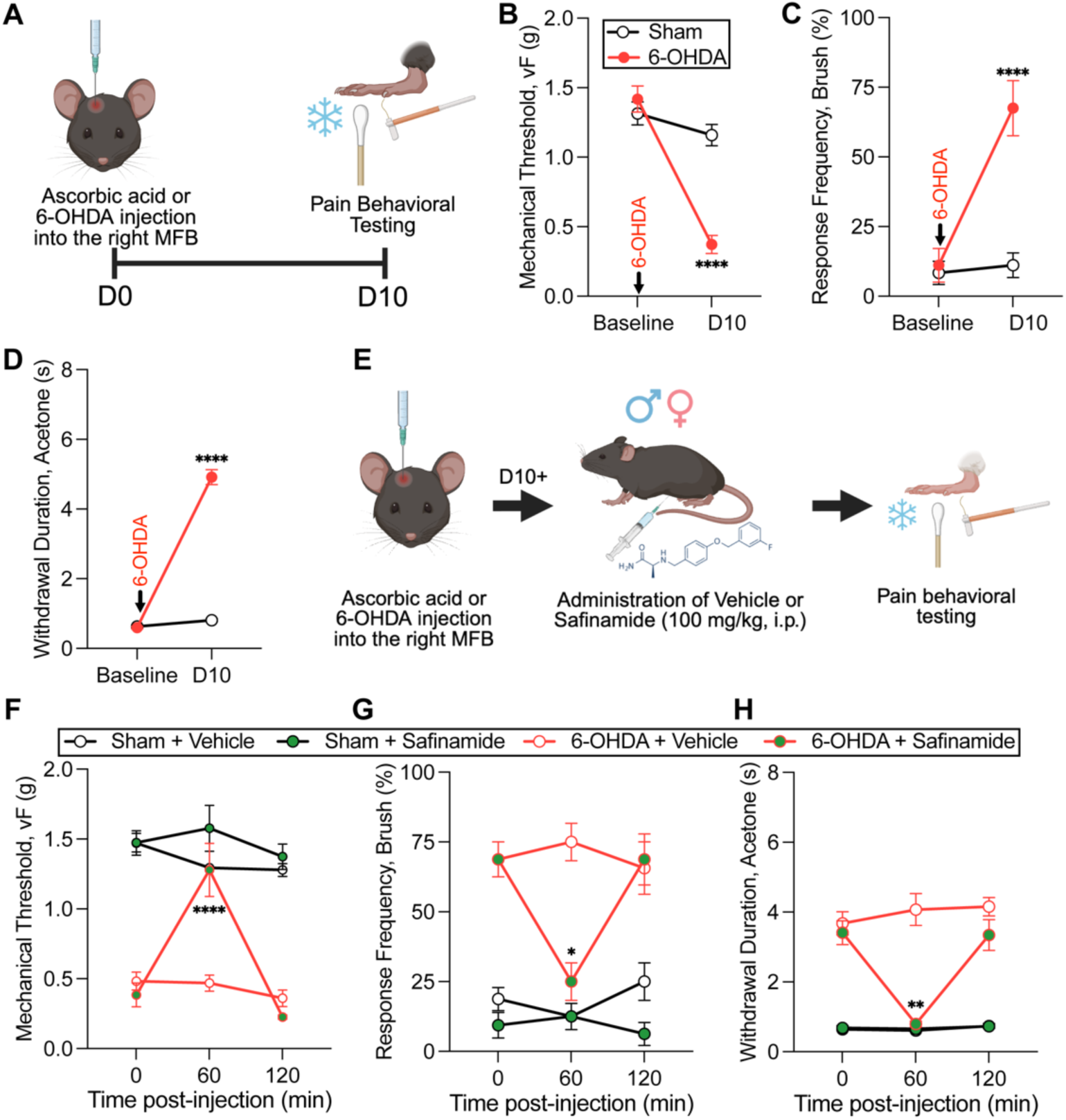
Safinamide reverses established pain-like behaviors in experimental Parkinson’s disease. **(A)** Experimental timeline for unilateral injection of ascorbic acid (sham) or 6-hydroxydopamine (6-OHDA) into the right medial forebrain bundle (MFB), followed by assessment of pain-like behaviors at baseline and 10 days (D10) post-lesion. **(B)** Mechanical withdrawal thresholds measured using von Frey (vF) filaments, **(C)** brush-evoked response frequency, and **(D)** acetone-evoked withdrawal duration at baseline and D10 in sham and 6-OHDA mice (n = 9-10 mice/group), demonstrating the development of mechanical hypersensitivity, dynamic mechanical allodynia, and cold hypersensitivity following 6-OHDA lesion. For **B–D**, statistical comparisons were performed using repeated-measures mixed-effects model followed by Holm-Šidák’s multiple comparisons test. **(E)** Experimental design for acute safinamide administration. Beginning ≥10 days following unilateral MFB injection, sham and 6-OHDA mice received a single intraperitoneal injection of vehicle (0.9% sodium chloride) or safinamide (100 mg/kg), followed by pain behavioral testing over 120 min. **(F)** Mechanical withdrawal thresholds, **(G)** brush-evoked response frequency, and **(H)** acetone-evoked withdrawal duration following vehicle or safinamide administration in sham + vehicle, sham + safinamide, 6-OHDA + vehicle, and 6-OHDA + safinamide mice (n = 8 mice/group). Safinamide rapidly reversed 6-OHDA-induced mechanical hypersensitivity, dynamic mechanical allodynia, and cold hypersensitivity, with significant effects observed at 60 min post-injection. For **F–H**, statistical comparisons were performed using repeated-measures three-way ANOVA followed by Tukey’s multiple comparisons test. Data are presented as mean ± SEM. * *P* < 0.05, ** *P* < 0.01, **** *P* < 0.0001.

### Parkinsonian neurodegeneration enhances NaV1.7-dependent sodium currents in primary sensory neurons

Having identified Na_V_1.7 as a target of safinamide, we next asked whether Na_V_1.7 function is altered in primary sensory neurons following nigrostriatal neurodegeneration. Whole-cell voltage-clamp recordings were performed in small-diameter DRG neurons isolated from sham- and 6-OHDA-treated mice in the absence or presence of 5 nM ProTx-II (**Fig. 4A**). Representative current traces revealed substantially larger inward sodium currents in neurons from 6-OHDA mice compared with sham controls, whereas currents recorded in the presence of ProTx-II were markedly reduced in both groups (**Fig. 4B**). Consistent with these recordings, 6-OHDA increased sodium current density across the current-voltage relationship (**Fig. 4C**) and increased peak sodium current density by approximately 38% compared with sham controls (**Fig. 4D**). ProTx-II reduced peak sodium current density by approximately 60% in sham neurons and 53% in 6-OHDA neurons and eliminated the significant difference in peak current between sham and 6-OHDA neurons (**Fig. 4, C and D**). These findings demonstrate that ProTx-II-sensitive Na_V_1.7 current contributes substantially to the increased sodium current observed following nigrostriatal neurodegeneration.

**Figure 4.**
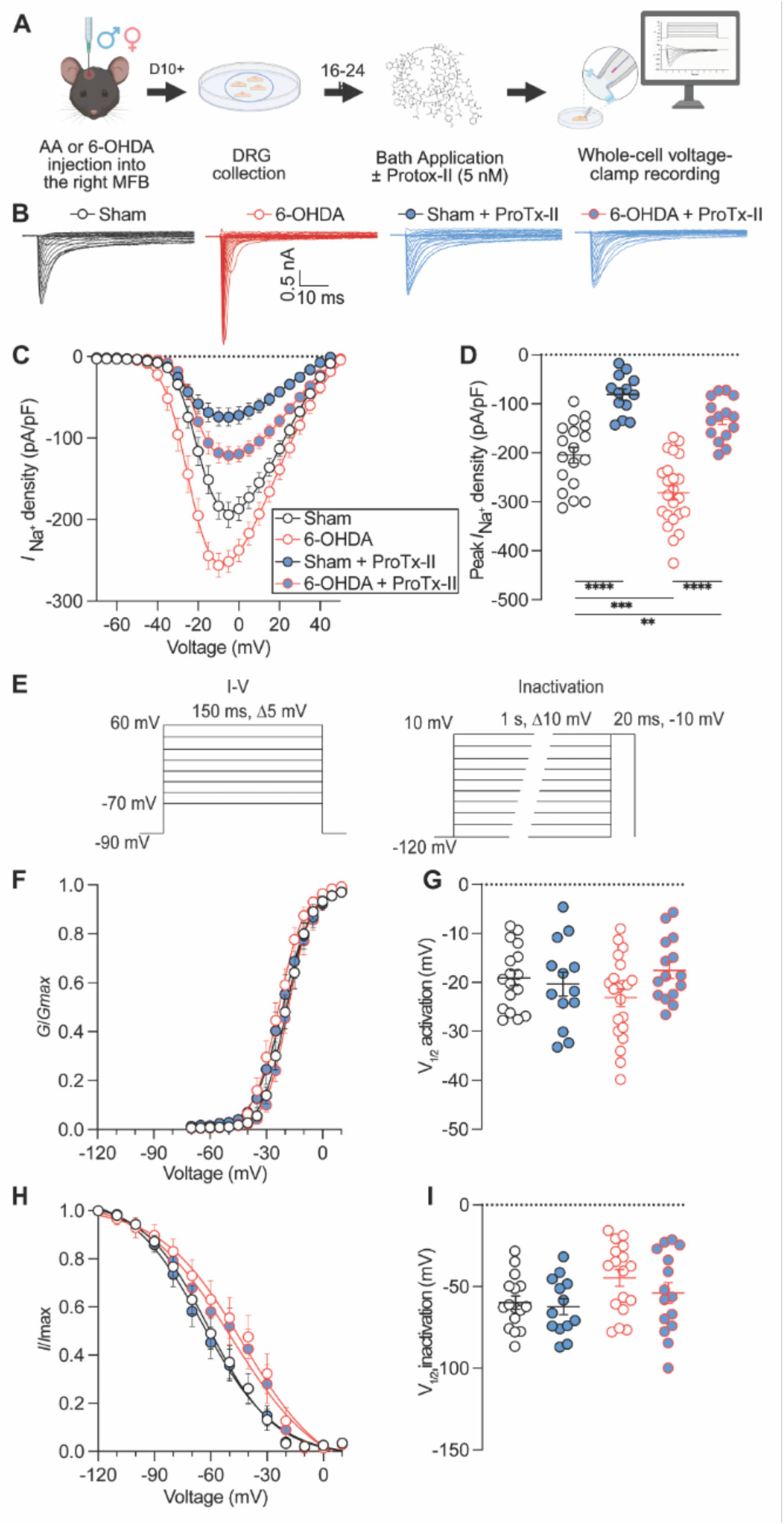
Nigrostriatal dopaminergic neurodegeneration increases Na_V_1.7-dependent sodium current in primary sensory neurons. **(A)** Experimental design for whole-cell voltage-clamp recordings from dorsal root ganglion (DRG) neurons isolated from male and female mice ≥10 days following unilateral injection of ascorbic acid (AA; sham) or 6-hydroxydopamine (6-OHDA) into the right medial forebrain bundle (MFB). Dissociated DRG neurons were cultured for 16–24 h and treated during recording with vehicle (0.1% DMSO) or ProTx-II (5 nM). **(B)** Representative whole-cell sodium current traces from sham, 6-OHDA, sham + ProTx-II, and 6-OHDA + ProTx-II groups. **(C)** Current–voltage (I–V) relationships for total voltage-gated sodium current in DRG neurons from sham and 6-OHDA mice in the absence or presence of ProTx-II. **(D)** Peak sodium current density for each experimental condition. DRG neurons from 6-OHDA mice exhibited increased peak sodium current density compared with sham controls, whereas ProTx-II reduced peak sodium current density and eliminated the difference between sham and 6-OHDA neurons. **(E)** Voltage-clamp protocols used to determine sodium current activation and steady-state inactivation. **(F)** Voltage dependence of activation and **(G)** half-maximal voltage of activation (V_1/2_ activation) derived from Boltzmann fits. **(H)** Voltage dependence of steady-state inactivation and **(I)** half-maximal voltage of inactivation (V_1/2_ inactivation) derived from Boltzmann fits. For **D, G, and I**, statistical comparisons were performed using two-way ANOVA. n = 13-23 cells/group. Data are presented as mean ± SEM with individual cells shown where applicable. **P < 0.01, ***P < 0.001, ****P < 0.0001.

We next examined whether dopaminergic neurodegeneration altered voltage-dependent sodium channel gating using activation and steady-state inactivation protocols (**Fig. 4E**). The voltage dependence of activation was unchanged across experimental groups (**Fig. 4F**), with no significant differences in V_1/2_ activation (**Fig. 4G**). Steady-state inactivation was also examined (**Fig. 4H**), and analysis of V_1/2_ inactivation revealed a significant effect of surgery (*F*(1,57) = 5.039, *P* = 0.0287; **Fig. 4I**). Thus, nigrostriatal neurodegeneration increases the Na_V_1.7-dependent contribution to sensory neuron sodium current and alters voltage-dependent channel gating. Together, these findings demonstrate that nigrostriatal neurodegeneration functionally remodels Na_V_1.7-mediated sodium currents in primary sensory neurons.

### Pharmacological disruption of CRMP2-dependent NaV1.7 regulation normalizes DRG neuron hyperexcitability following Parkinsonian neurodegeneration

Having identified increased Na_V_1.7 activity following nigrostriatal neurodegeneration, we next asked whether targeting CRMP2-dependent regulation of Na_V_1.7 could normalize the resulting sensory neuron hyperexcitability. Small-diameter DRG neurons were isolated from sham- and 6-OHDA-treated mice and incubated with vehicle or Compound 194 (C194; 5 μM), a small molecule that disrupts CRMP2-dependent regulation of Na_V_1.7 by preventing CRMP2 SUMOylation (36), prior to whole-cell current-clamp recording (**Fig. 5A**). Representative traces demonstrated increased repetitive action potential firing in neurons from 6-OHDA mice, whereas C194 markedly reduced firing in neurons from 6-OHDA mice (**Fig. 5B**).

**Figure 5.**
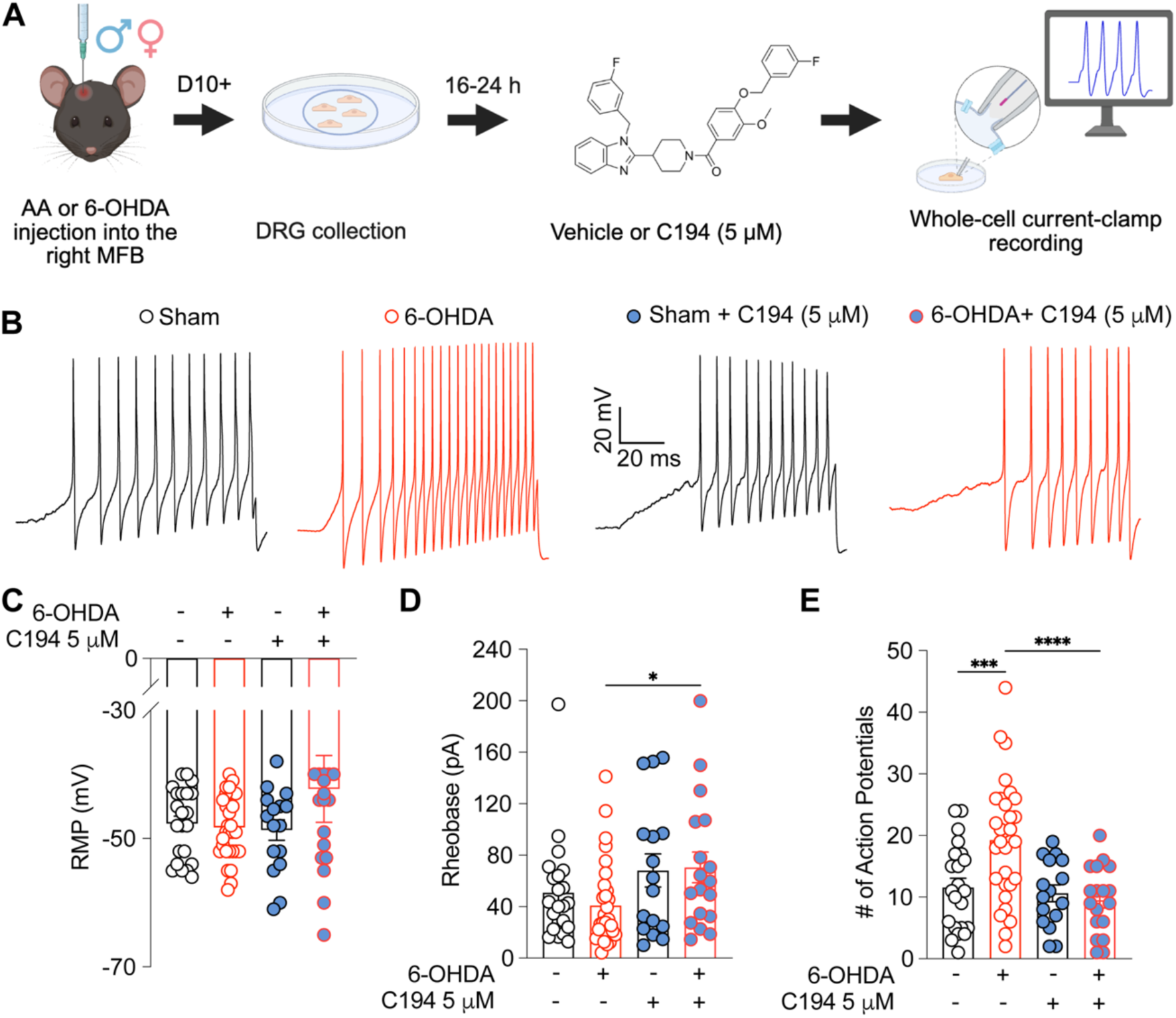
Pharmacological antagonism of CRMP2-dependent Na_V_1.7 regulation normalizes DRG neuron hyperexcitability following Parkinsonian neurodegeneration. **(A)** Experimental design for whole-cell current-clamp recordings. Dorsal root ganglia (DRG) were collected from male and female mice ≥10 days following unilateral injection of ascorbic acid (AA; sham) or 6-hydroxydopamine (6-OHDA) into the right medial forebrain bundle (MFB). Dissociated DRG neurons were incubated for 16–24 h with vehicle (0.1% DMSO) or Compound 194 (C194; 5 μM) prior to electrophysiological recording. **(B)** Representative current-clamp traces from small-diameter DRG neurons isolated from Sham, 6-OHDA, Sham + C194, and 6-OHDA + C194 mice during a ramp depolarizing current injection. **(C)** Resting membrane potential (RMP) of DRG neurons from Sham, 6-OHDA, Sham + C194, and 6-OHDA + C194 groups (n = 16-30 cells /group). **(D)** Rheobase of DRG neurons from Sham, 6-OHDA, Sham + C194, and 6-OHDA + C194 groups (n = 16-30 cells /group). C194 significantly increased the current required to initiate action potential firing. **(E)** Number of action potentials generated in response to depolarizing current injection in Sham, 6-OHDA, Sham + C194, and 6-OHDA + C194 groups (n = 16-30 cells /group). DRG neurons from 6-OHDA mice exhibited increased repetitive firing compared with sham controls, which was normalized by C194 treatment. For **C–E**, statistical comparisons were performed using two-way ANOVA followed by uncorrected Fisher’s LSD. Data are presented as mean ± SEM with individual cells shown. * *P* < 0.05, *** *P* < 0.001, **** *P* < 0.0001.

Resting membrane potential was not significantly altered by 6-OHDA lesioning or C194 treatment (**Fig. 5C**), indicating that the hyperexcitable phenotype was not attributable to a sustained change in baseline membrane potential. In contrast, C194 significantly increased rheobase in neurons from 6-OHDA mice, increasing the amount of current required to elicit an action potential (**Fig. 5D**). Consistent with this effect, DRG neurons from 6-OHDA mice generated significantly more action potentials in response to depolarizing current injection than sham neurons, and C194 normalized this increased repetitive firing (**Fig. 5E**). Together, these findings demonstrate that pharmacological antagonism of CRMP2-dependent Na_V_1.7 regulation normalizes the altered intrinsic excitability of primary sensory neurons following nigrostriatal neurodegeneration.

### Pharmacological disruption of CRMP2-dependent NaV1.7 regulation reverses established Parkinsonian pain-like behavior

The ability of C194 to normalize 6-OHDA-mediated DRG neuron hyperexcitability suggested that targeting CRMP2-dependent Na_V_1.7 regulation could also suppress established Parkinsonian pain. To test this, mice received a single intraperitoneal injection of vehicle or C194 (10 mg/kg) (36), and pain-like behaviors were assessed one hour after administration (**Fig. 6A**). C194 completely reversed mechanical hypersensitivity in 6-OHDA mice, restoring withdrawal thresholds to sham levels (**Fig. 6B**). C194 also abolished dynamic mechanical allodynia and markedly reduced cold hypersensitivity (**Fig. 6, C and D**). In addition, C194 significantly increased withdrawal latency in the 52°C hot plate assay (**Fig. 6E**). Together, these findings demonstrate that acute pharmacological disruption of CRMP2-dependent Na_V_1.7 regulation reverses established multimodal pain-like behavior following nigrostriatal neurodegeneration.

**Figure 6.**
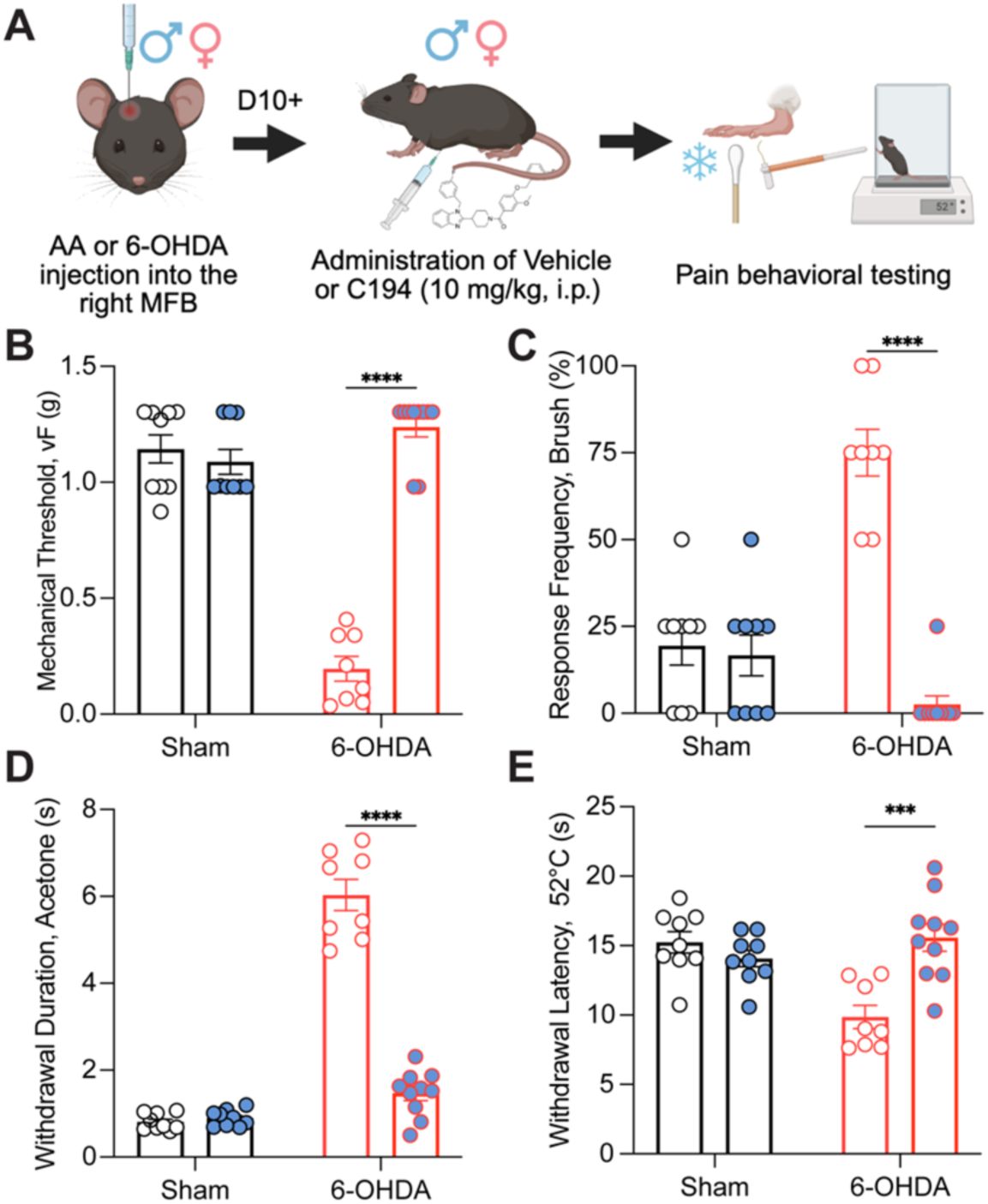
Pharmacological disruption of CRMP2-dependent Na_V_1.7 regulation reverses Parkinsonian pain-like behavior. **(A)** Experimental design for acute Compound 194 (C194) administration. Beginning ≥10 days following unilateral injection of ascorbic acid (AA; sham) or 6-hydroxydopamine (6-OHDA) into the right medial forebrain bundle (MFB), male and female mice received a single intraperitoneal injection of vehicle (1:1:8; DMSO, Tween 80, 0.9% sodium chloride) or C194 (10 mg/kg), followed by pain behavioral testing 1 h after administration. **(B)** Mechanical withdrawal thresholds measured using von Frey (vF) filaments, **(C)** brush-evoked response frequency, **(D)** acetone-evoked withdrawal duration, and **(E)** withdrawal latency in the 52°C hot plate assay in sham and 6-OHDA mice following vehicle or C194 administration (n = 8-10 mice/group). C194 reversed 6-OHDA-induced mechanical hypersensitivity, dynamic mechanical allodynia, and cold hypersensitivity and increased heat withdrawal latency. For **B–E**, statistical comparisons were performed using two-way ANOVA followed by Uncorrected Fisher’s LSD. Data are presented as mean ± SEM. *** *P* < 0.001, **** *P* < 0.0001.

### Repeated C194 administration produces sustained analgesic efficacy throughout Parkinsonian disease progression

To determine whether the analgesic effects of C194 are maintained with repeated administration, 6-OHDA mice received C194 (10 mg/kg, i.p.) at 2-, 4-, 6-, and 12-weeks post-lesion. Sensory behaviors were assessed immediately before treatment and 2 and 24 hours after each administration (**Fig. 7A**). At each treatment time point, C194 rapidly increased mechanical withdrawal thresholds at 2 hours, with responses returning toward pretreatment levels by 24 hours (**Fig. 7B**). Importantly, the magnitude of the C194-induced increase in mechanical withdrawal threshold was comparable across the four treatment sessions (**Fig. 7C**). C194 similarly reduced brush-evoked responses following each administration (**Fig. 7D**), with no detectable reduction in the magnitude of this effect across 2, 4, 6, and 12 weeks (**Fig. 7E**). Acetone-evoked withdrawal duration was likewise reduced 2 hours after each C194 treatment and returned toward pretreatment levels by 24 hours (**Fig. 7F**), while the magnitude of the reduction in cold hypersensitivity remained consistent across repeated administrations (**Fig. 7G**). Together, these findings demonstrate that C194 repeatedly reverses established mechanical, dynamic mechanical, and cold hypersensitivity over a 12-week period, with no detectable loss of analgesic efficacy across successive administrations.

**Figure 7.**
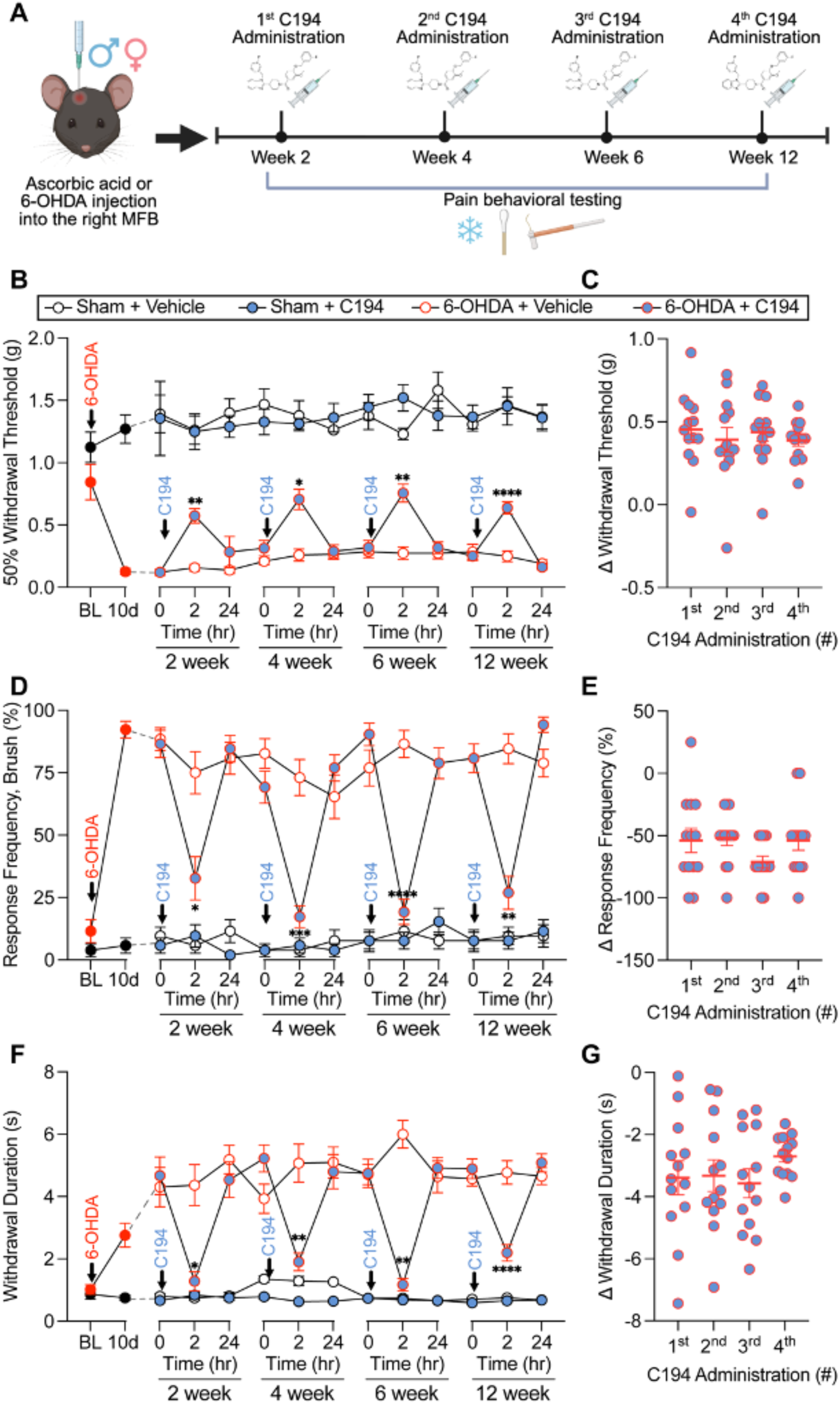
Repeated pharmacological antagonism of CRMP2-dependent Na_V_1.7 regulation produces sustained reversal of Parkinsonian pain-like behavior. **(A)** Experimental design for repeated Compound 194 (C194) administration. Following unilateral injection of ascorbic acid (sham) or 6-hydroxydopamine (6-OHDA) into the right medial forebrain bundle (MFB), male and female mice received vehicle (1:1:8; DMSO, Tween 80, 0.9% sodium chloride) or C194 (10 mg/kg, i.p.) at 2-, 4-, 6-, and 12-weeks post-lesion, with pain behavioral testing performed before and for 24 h following each administration. **(B)** Mechanical withdrawal thresholds measured using von Frey (vF) filaments and **(C)** change in mechanical withdrawal threshold following each C194 administration. **(D)** Brush-evoked response frequency and **(E)** change in brush-evoked response frequency following each C194 administration. **(F)** Acetone-evoked withdrawal duration and **(G)** change in acetone-evoked withdrawal duration following each C194 administration (n = 13 mice/group). Repeated C194 administration consistently reversed 6-OHDA-induced mechanical hypersensitivity, dynamic mechanical allodynia, and cold hypersensitivity across the 12-week observation period, with sensory responses returning toward pre-treatment levels by 24 h. For **B, D,** and **F**, statistical comparisons were performed using three-way repeated measures ANOVA followed by Tukey’s multiple-comparisons test. For **C, E,** and **G**, statistical comparisons were performed using one-way ANOVA. Data are preented as mean ± SEM. * *P* < 0.05, ** *P* < 0.01, *** *P* < 0.001, **** *P* < 0.0001.

### Intranasal C194 reverses established Parkinsonian pain-like behavior

To determine whether C194 retains analgesic efficacy through a noninvasive route of administration, mice with established 6-OHDA-induced hypersensitivity received a single intranasal administration of vehicle or C194 (40 mg/kg), followed by behavioral testing (**Supplementary Fig. S2A**). Intranasal C194 significantly increased mechanical withdrawal thresholds in 6-OHDA mice (**Supplementary Fig. S2B**) and reduced brush-evoked responses (**Supplementary Fig. S2C**). Intranasal C194 also markedly reduced acetone-evoked withdrawal duration, demonstrating efficacy against cold hypersensitivity (**Supplementary Fig. S2D**). Together, these findings demonstrate that C194 reverses multiple modalities of established Parkinsonian pain-like behavior following intranasal administration and support the feasibility of noninvasive delivery for targeting CRMP2-dependent Na_V_1.7 regulation.

### Genetic disruption of CRMP2-dependent NaV1.7 regulation prevents the development of Parkinsonian pain

Our pharmacological studies demonstrated that disrupting CRMP2-dependent Na_V_1.7 regulation reverses established pain-like behaviors following Parkinsonian neurodegeneration. We next sought to determine whether CRMP2-dependent regulation of Na_V_1.7 is necessary for the development of Parkinsonian pain using a complementary genetic approach. CRMP2 interacts with a unique 15-amino-acid CRMP2 regulatory sequence (CRS; residues 706–720, SRQKCPPWWYRFAHK) located within the first intracellular loop of Na_V_1.7, which is required for CRMP2-dependent regulation of channel trafficking and function (38). To determine whether CRS is required for Parkinsonian pain, we generated mice carrying a modified *Scn9a* allele in which the endogenous CRS was replaced by a FLAG epitope sequence. Homozygous Na_V_1.7^CRSΔ/Δ^ (ΔCRS) mice therefore lack an intact CRS on either Na_V_1.7 allele, disrupting CRS-dependent regulation of the channel (**Fig. 8A**).

**Figure 8.**
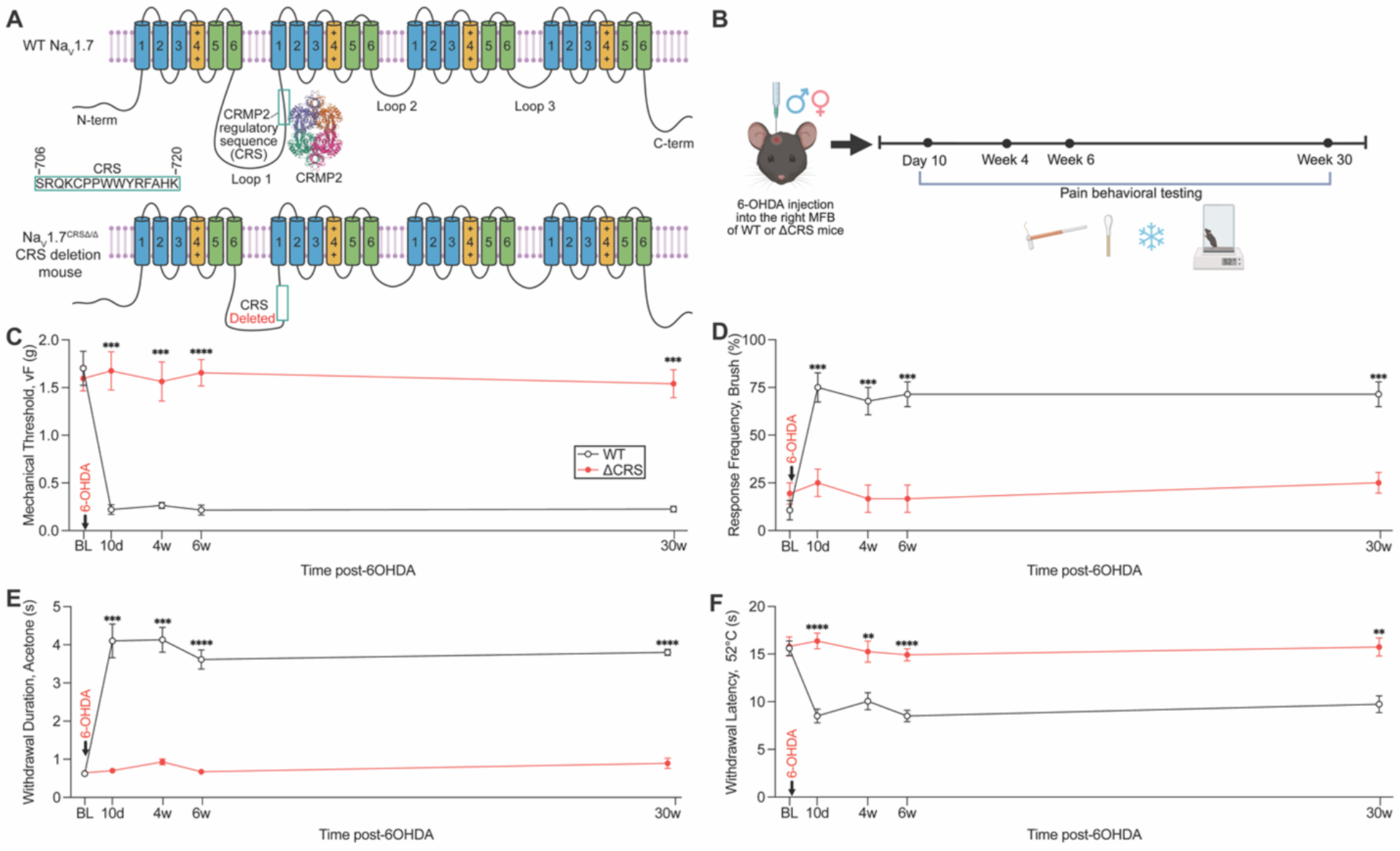
Genetic disruption of CRMP2-dependent Na_V_1.7 regulation prevents the development of Parkinsonian pain-like behavior. **A)** Schematic of wild-type (WT) Na_V_1.7 and the Na_V_1.7^CRSΔ/Δ^ (ΔCRS) mouse, in which the endogenous CRMP2 regulatory sequence (CRS; residues 706–720) within the first intracellular loop of Na_V_1.7 is replaced by a FLAG epitope sequence, disrupting CRS-dependent regulation of the channel. **(B)** Experimental design for longitudinal assessment of pain-like behaviors following unilateral injection of 6-hydroxydopamine (6-OHDA) into the right medial forebrain bundle (MFB) of WT and ΔCRS mice. Pain behavioral testing was performed at baseline and 10-days, 4-, 6-, and 30-weeks post-lesion. **(C)** Mechanical withdrawal thresholds measured using von Frey (vF) filaments, **(D)** brush-evoked response frequency, **(E)** acetone-evoked withdrawal duration, and **(F)** withdrawal latency in the 52°C hot plate assay in WT and ΔCRS mice before and following 6-OHDA lesion (n = 7-9 mice/group). WT mice developed persistent mechanical hypersensitivity, dynamic mechanical allodynia, cold hypersensitivity, and heat hypersensitivity following 6-OHDA lesion, whereas ΔCRS mice were protected from the development of pain-like behaviors throughout the 30-week observation period. For **C–F**, statistical comparisons were performed using two-way repeated-measures mixed-effects analysis followed by Holm-Šidák’s multiple comparisons test. Data are presented as mean ± SEM. ** *P* < 0.01, *** *P* < 0.001, **** *P* < 0.0001.

We first characterized the functional consequences of CRS disruption in naïve ΔCRS mice. Whole-cell voltage-clamp recordings from primary sensory neurons demonstrated a marked reduction in total sodium current density, with peak sodium current reduced by approximately 54% in ΔCRS neurons compared with wild-type neurons (**Supplementary Fig. S3, B and C**). CRS disruption also significantly altered the voltage dependence of channel activation but not steady-state inactivation (**Supplementary Fig. S3, D–F**), indicating that disruption of the NaV1.7 CRMP2 regulatory sequence affects not only sodium current magnitude but also channel gating. Despite these electrophysiological changes, ΔCRS mice exhibited largely preserved baseline somatosensory function, including mechanical, cold, and pin-prick responses, with only modest differences in thermal withdrawal at higher temperatures (**Supplementary Fig. S4**). We next challenged these mice using spared nerve injury (SNI), a chronic neuropathic pain model in which Na_V_1.7 contributes strongly to the development of hypersensitivity. Whereas wild-type mice developed robust and persistent mechanical and cold hypersensitivity following SNI, ΔCRS mice were markedly resistant to the development of neuropathic pain-like behavior in both males and females (**Supplementary Fig. S5**). Together, these findings demonstrate that disruption of the CRS produces substantial functional remodeling of sensory neuron sodium currents and strongly attenuates pathological pain while largely preserving baseline nociceptive function.

Having established that pharmacological disruption of CRMP2-dependent Na_V_1.7 regulation reverses established pain, we next asked whether an intact Na_V_1.7 CRS is required for the development of Parkinsonian pain. WT and ΔCRS mice received unilateral 6-OHDA injections into the right MFB and were followed longitudinally for up to 30 weeks (**Fig. 8, A and B**). Importantly, CRS disruption did not protect against the motor consequences of nigrostriatal dopaminergic neurodegeneration. Following 6-OHDA lesioning, WT and ΔCRS mice exhibited comparable apomorphine-induced contralateral rotations, rotarod performance, and nest-building deficits (**Supplementary Fig. S6, A-E**), indicating that disruption of CRMP2-dependent Na_V_1.7 regulation did not attenuate the underlying Parkinsonian phenotype. Despite comparable motor impairment, the sensory consequences of 6-OHDA lesioning were strikingly different between genotypes. WT mice developed robust mechanical hypersensitivity, dynamic mechanical allodynia, cold hypersensitivity, and heat hypersensitivity that persisted throughout the 30-week observation period (**Fig. 8, C-F**). In contrast, ΔCRS mice were protected from the development of these pain-like behaviors, with sensory responses remaining near baseline across the entire observation period (**Fig. 8, C-F**). Thus, genetic disruption of the Na_V_1.7 CRS dissociates the sensory consequences of nigrostriatal neurodegeneration from the underlying Parkinsonian motor phenotype and prevents the development of persistent pain for at least 30 weeks.

### Human Parkinson’s disease DRG exhibit selective transcriptional features associated with sensory neuron excitability

To determine whether human PD is associated with molecular alterations within peripheral sensory ganglia, we performed bulk RNA sequencing of thoracic DRG obtained from individuals with PD (n = 8), age-matched controls (n = 8), and young controls (n = 4) (**Fig. 9A** and **Supplementary Table S2**). Global expression distributions were comparable across all 20 donors, and pairwise transcriptomic correlations were uniformly high, with no individual sample exhibiting a markedly divergent global expression profile (**Supplementary Figs. S7** and **S8**). Principal component analysis of the 2,000 most variable genes demonstrated substantial overlap between PD and age-matched control samples, whereas the young control samples provided an additional reference for age-associated variation (**Fig. 9B**). We first performed an exploratory genome-wide comparison of PD and age-matched control DRG using an age-adjusted limma model. Consistent with the substantial overlap observed by PCA, relatively limited transcriptional differences were identified between groups (**Supplementary Fig. S9A**). The 30 genes with the smallest nominal P values were visualized across individual donors, revealing substantial interindividual heterogeneity without clear separation between PD and age-matched control donors (**Supplementary Fig. S9B**). Together, these findings indicate relatively limited global transcriptional remodeling of the DRG in PD.

**Figure 9.**
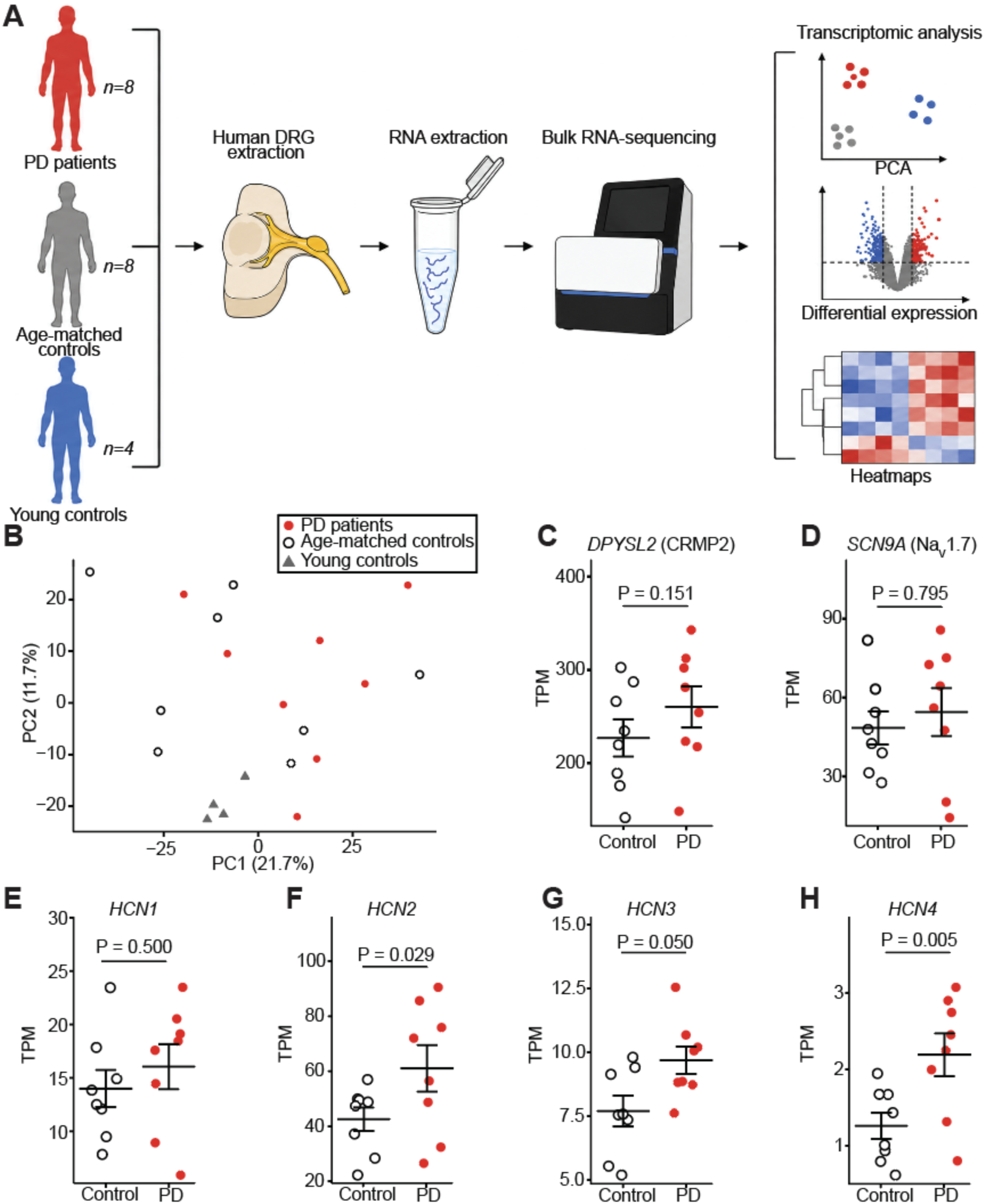
Human Parkinson’s disease DRG exhibit selective transcriptional features associated with sensory neuron excitability. associated with sensory neuron excitability. **(A)** Experimental workflow for bulk RNA sequencing of human dorsal root ganglia (DRG) from individuals with Parkinson’s disease (PD; n = 8), age-matched controls (n = 8), and young controls (n = 4). **(B)** Principal component analysis of the 2,000 most variable genes across all donors. **(C-D)** Transcript abundance of *DPYSL2*, encoding CRMP2 (C), and *SCN9A*, encoding Na_V_1.7 (D), in PD and age-matched control DRG. **(E-H)** Transcript abundance of *HCN1* **(E)**, *HCN2* **(F)**, *HCN3* **(G)**, and *HCN4* **(H)** in PD and age-matched control DRG. Differential expression between PD and age-matched controls was assessed using limma with age included as a covariate and empirical Bayes moderation. For **C-H**, individual points represent individual donor TPM and are shown with mean ± SEM; exact *P* values displayed above each plot were derived from the age-adjusted limma model.

We next examined genes directly related to the sensory neuron mechanisms identified in our mouse studies. *DPYSL2*, encoding CRMP2, was unchanged in PD DRG compared with age-matched controls (*P* = 0.151; **Fig. 9C**), as was *SCN9A*, encoding Na_V_1.7 (*P* = 0.795; **Fig. 9D**). Given our finding of increased Na_V_1.7-dependent current in Parkinsonian sensory neurons, we next examined HCN channels, as HCN-mediated *I*_h_ has been shown to enhance Na_V_1.7 inactivation, reduce Na_V_1.7-mediated sodium influx, and suppress Na_V_1.7-dependent DRG neuron hyperexcitability (43). *HCN1* expression was unchanged in PD DRG (*P* = 0.500; **Fig. 9E**), whereas *HCN2* showed higher expression (*P* = 0.029; **Fig. 9F**). *HCN3* showed a similar pattern (*P* = 0.050; **Fig. 9G**), and *HCN4* expression was also higher in PD DRG (*P* = 0.005; **Fig. 9H**). Together, these findings identify altered expression patterns across multiple HCN channel family members in human PD DRG despite unchanged *SCN9A* and *DPYSL2* transcript abundance.

Finally, the young control cohort allowed us to examine whether transcriptional patterns across sensory signaling pathways were associated with normal aging. Three-group heatmaps were generated for ion-channel and intrinsic excitability, neurotrophic and pain-signaling, immune and inflammatory, and nerve injury and regeneration gene sets across young controls, age-matched controls, and PD donors (**Supplementary Figs. S10-S13**). These analyses revealed substantial gene- and donor-specific variability and did not show a uniform progression from young control to age-matched control to PD across the examined pathways.

Collectively, these findings demonstrate relatively limited global transcriptional remodeling in human PD DRG, with no transcriptional induction of *SCN9A* or *DPYSL2* despite the functional Na_V_1.7 phenotype identified in mice. In contrast, expression patterns across multiple HCN channel family members differed in PD DRG, identifying HCN channels as candidates for future functional investigation of altered peripheral sensory neuron excitability in PD.

## Methods

### Sex as a Biological Variable

Both male and female mice were included with approximately equal representation. Experiments were not powered to detect sex-specific effects; therefore, sexes were pooled for the primary analyses.

### Animals

Adult C57BL/6J (Jackson Laboratory, # 000664), and Na_V_1.7^CRSΔ/Δ^ (ΔCRS) male and female mice were group housed, provided access to food and water *ad libitum*, and maintained on a 12:12 hour light:dark cycle in temperature and humidity-controlled rooms. Adult mice between 8 weeks and 10 months of age were used for all experiments. All experimental procedures were approved by the Institutional Animal Care and Use Committees of University of Florida (IACUC202400000002) and were performed in accordance with the National Institutes of Health Guide for the Care and Use of Laboratory Animals.

### Generation and genotyping of NaV1.7 ΔCRS mutant mice

Na_V_1.7 CRMP2 regulatory sequence (CRS) mutant mice were generated by targeted modification of the endogenous *Scn9a* locus. The 15-amino-acid sequence encoding the Na_V_1.7 CRS within the first intracellular loop of the channel was deleted and replaced with a FLAG epitope sequence, thereby disrupting the native CRS while maintaining the surrounding *Scn9a* coding sequence. The modified allele was initially generated and maintained as a heterozygous line and subsequently bred to homozygosity. Homozygous mice, referred to here as Na_V_1.7^CRSΔ/Δ^ (ΔCRS), therefore express Na_V_1.7 in which both endogenous CRS alleles are replaced by the FLAG-containing modified sequence. Initial genotyping was performed by PCR using primers flanking the modified region (forward, 5′-GATTTTGTTGCCGATTTCTACC-3′; reverse, 5′- AAGGTGTTTAAAACTATGCAAATGG-3′). Because the size difference between the wild-type and modified PCR products was small, genotype was resolved by BbsI restriction digestion using a restriction site introduced within the FLAG sequence. Wild-type alleles produced an undigested 262-bp product, whereas the modified allele produced 101- and 143-bp restriction fragments. Accordingly, heterozygous mice displayed all three fragments, whereas homozygous ΔCRS mice displayed only the 101- and 143-bp products. An additional FLAG-specific forward primer (5′- GAAGACTACAAAGACGATGACG-3′) paired with the reverse screening primer generated a 146-bp product specific to the modified allele. All ΔCRS mice used in this study were genotyped by Transnetyx following collection of ear-punch tissue. Genotyping was performed to distinguish wild-type, heterozygous, and homozygous CRS-modified alleles, and genotype was confirmed for each experimental animal prior to inclusion in the study. Only mice with a confirmed genotype were included in experimental analyses.

### Randomization and Blinding

Animals were randomly assigned to experimental groups using simple randomization. Behavioral testing, image quantification, and electrophysiological data analysis were conducted by investigators blinded to treatment groups whenever feasible.

### Sample Size

No formal *a priori* power analysis was conducted. Sample sizes were determined based on prior studies from our laboratory and published literature that employed comparable behavioral, histological, and electrophysiological measures. Sample sizes for individual experiments are provided in the corresponding figure legends.

### Drug Administration

Safinamide (MedChemExpress, Cat#HY-70057A; 100 mg/kg in 0.9% sodium chloride, i.p.) and vehicle (0.9% sodium chloride, i.p.) were injected fresh each day using a 30-gauge needle while mice were briefly scruffed. Compound 194 (Source: National Center for Advancing Translational Sciences; 10 mg/kg in 10:10:80, DMSO, Tween-80, 0.9% sodium chloride, i.p.) and vehicle (10:10:80, DMSO, Tween-80, 0.9% sodium chloride, i.p.) were freshly prepared each day and administered with a 30-gauge needle while mice were scruffed briefly. Additionally, C194 ((4- (1-(3-fluorobenzyl)-1H-benzo[d]imidazol-2-yl)piperidin-1-yl)(4-((3-fluorobenzyl)oxy)-3-methoxyphenyl)methanone) (36) was administered intranasally at a dose of 1 mg/mouse (∼40 mg/kg) in a total volume of 20 µL of 100% DMSO, with 10 µL delivered into each nostril. Vehicle-treated mice received an equivalent volume of 100% DMSO. Intranasal administration was performed using a micropipette while mice were lightly anesthetized with approximately 1.5% isoflurane.

### Surgery

#### Unilateral 6-hydroxydopamine (6-OHDA) surgery

Surgeries were conducted as previously described (41). In brief, mice were injected with desipramine hydrochloride (Sigma-Aldrich, Cat#D3900, 25 mg/kg, i.p.) 30 minutes before surgery to protect noradrenergic neurons. Mice were anesthetized with isoflurane (1.5–2.5%) and secured in a stereotaxic apparatus. The scalp was shaved and sterilized with alternating washes of 70% ethanol and povidone-iodine before a midline incision approximately 1 cm in length was made. A small craniotomy was drilled over the right medial forebrain bundle (MFB; coordinates relative to bregma: AP -1.0 mm, ML -1.2 mm, DV -4.8 mm from the surface of the brain), and 0.3 μL of 6-OHDA (Tocris, Cat#2547, 12 μg/μL in 0.2% ascorbic acid) was injected at a rate of 0.06 μL/min using a Nanoject III Auto-Nanoliter Injector (Drummond). Sham-operated mice received an equivalent volume of vehicle (0.2% ascorbic acid) at the same stereotaxic coordinates. Following injection, the injector was left in place for an additional 5 minutes to facilitate diffusion before being slowly withdrawn. The incision was closed with 6-0 nylon monofilament sutures, and mice received a single postoperative injection of meloxicam (20 mg/kg, s.c.). Animals recovered on a heating pad before being returned to their home cages. Postoperative analgesia was continued for 72 hours using acetaminophen-supplemented drinking water. Additional supportive care was provided as needed for up to 14 days following 6-OHDA lesion and included DietGel (Clear H2O, Cat#72-06-5022), STAT High Calorie Liquid Diet (PRN, Cat#30022409, p.o.), and supplemental subcutaneous saline for hydration. Supplemental heat was provided as needed by positioning approximately half of the cage over a heating pad, allowing animals to move away from the heat source. Male mice were additionally monitored for penile prolapse and accumulation of urinary crystals. When present, urinary crystals were gently removed, Surgilube was applied to maintain tissue lubrication, and the prolapsed penis was manually reduced into the prepuce. Animals were subsequently monitored to confirm successful reduction and normal urination.

#### Spared nerve injury

Mice were anesthetized with isoflurane (1.5–2.5%), and the left hindlimb was shaved and sterilized with alternating washes of 70% ethanol and povidone-iodine. An incision was made along the lateral surface of the left thigh, and the biceps femoris muscle was bluntly dissected to expose the three terminal branches of the sciatic nerve. The common peroneal and tibial nerves were tightly ligated with 6-0 silk suture and transected approximately 2 mm distal to the ligation, while the sural nerve was left intact. The incision was closed in two layers: the muscle was reapproximated with 5-0 absorbable suture, and the skin was closed with 9-mm wound clips. Animals were allowed to recover for 7 days before behavioral testing.

### Behavioral Testing

#### Static Mechanical Sensitivity (von Frey)

Punctate mechanical sensitivity was assessed using a series of calibrated von Frey monofilaments (Braintree Scientific, Cat# 58011; 0.007–6.0 g). Prior to testing, mice were habituated for 60 minutes in individual plexiglass chambers (80 × 80 × 110 mm) positioned above an elevated wire mesh surface. Mice were then tested using the 0.4g filament, which was applied perpendicular to the lateral plantar surface of the left hindpaw with sufficient pressure to produce slight bending of the filament. A rapid paw withdrawal occurring within 4 seconds was scored as a positive response, after which the next lower-force filament was applied. If no withdrawal occurred, the next higher-force filament was used. Mechanical withdrawal thresholds were calculated using the up-down method to determine 50% withdrawal threshold for each mouse.

#### Dynamic Mechanical Sensitivity (Brush)

Dynamic tactile sensitivity was evaluated following von Frey testing in the same chambers. A cotton swab was teased to expand to approximately three times its original size and then lightly brushed across the left hindpaw from the heel towards the toes. A rapid withdrawal of the paw was considered a positive response. Four stimulation trials were performed for each mouse, and response frequency was calculated as the percentage of trials resulting in a paw withdrawal.

#### Mechanical Hyperalgesia (Pin-Prick)

Noxious mechanical sensitivity was assessed by gently applying a blunted pin to the plantar surface of the hindpaw without penetrating the skin. A rapid paw withdrawal was scored as a positive response. Four trials were performed per animal and response frequency was calculated as the percentage of trials eliciting withdrawal.

#### Cold Sensitivity (Acetone)

Responses to innocuous cold were assessed by applying approximately 10 μL of acetone (VWR, Cat# BDH1101-1LP) to the plantar surface of the left hindpaw. Acetone was administered using a syringe connected to PE-90 tubing with a flared tip measuring approximately 3.5 mm. Testing was performed immediately after the brush assay while mice remained in the same chambers. Following acetone application, the cumulative duration of paw lifting, shaking, or licking was recorded over a 30 second observation period. Three trials were conducted for each mouse, and responses were averaged across trials.

#### Thermal Withdrawal Latency (Hot Plate)

Acute thermal nociception was assessed using a hot/cold plate apparatus (Ugo Basile; Stoelting, Cat# 55075) maintained at 52.0°C. Mice were acclimated to the behavioral testing room for a minimum of 30 minutes before testing. For each trial, a mouse was placed on the apparatus and the latency to nocifensive response, defined as paw licking, lifting, or jumping, was recorded. If no response occurred, the trial was terminated after 30 seconds. Three trials were conducted per mouse with an intertrial interval of at least 10 minutes, and the mean withdrawal latency was calculated for subsequent statistical analysis. For baseline characterization of ΔCRS mice, thermal withdrawal latency was additionally assessed at 48°C and 55°C using the same apparatus.

#### Nesting Behavior

Nesting behavior was evaluated approximately 4 weeks following 6-OHDA lesioning as a measure of goal-directed and motivated behavior. Mice were individually housed in clean cages containing a fresh 10 × 10 cm nestlet for 12 hours during the dark cycle. Following the testing period, nests were scored using a six-point scale ranging from 0, indicating an untouched nestlet, to 5, indicating a well-formed, enclosed nest.

#### Motor Coordination and Balance (Accelerating Rotarod)

Motor coordination and balance were evaluated following 6-OHDA lesioning using an accelerating rotarod apparatus (IITC Life Science, Inc., Woodland Hills, CA, USA) consisting of five individual lanes with a 1.25-inch-diameter rotating rod. Mice were placed on the stationary rod before acceleration was initiated from 4 to 40 rpm over 300 s. Five trials were performed per mouse, and each trial ended when the mouse fell from the rod. Both latency to fall (s) and rotational speed at the time of falling (rpm) were recorded.

##### Apomorphine-Induced Rotation Test

Successful unilateral dopamine lesioning following injection with 6-OHDA was confirmed using an apomorphine-induced rotation test. Mice received an injection of apomorphine hydrochloride (Sigma-Aldrich, Cat#PHR2621, 0.25 mg/kg in 0.1% ascorbic acid, 0.9% saline, s.c.) or vehicle (0.1% ascorbic acid, 0.9% saline, s.c.) and were placed individually in a tall, cylindrical behavioral chamber (30 cm diameter) with an overhead camera. Full-body rotations were recorded for 45 minutes using AnyMaze software (Stoelting Co.), which tracked the head, mid-body, and tail of each animal. The number of both ipsilateral and contralateral rotations were recorded.

### Immunohistochemistry

At least 3 weeks following 6-OHDA administration, mice were deeply anesthetized and transcardially perfused using 30 mL ice-cold PBS followed by 30 mL ice-cold 10% neutral buffered formalin. The brain was extracted and post-fixed in 10% neutral buffered formalin overnight at 4°C. The brain was then cryoprotected in 30% sucrose for 5 days at 4°C, and three to five 35 µm sections containing the striatum were obtained from each animal using a cryostat. Tissue was washed three times in 1x phosphate buffered saline (PBS) and then incubated in blocking buffer (5% normal goat serum, 0.1% Triton X-100 in 1x PBS) for 90 minutes. Tissue was then incubated overnight in primary antibody (Rabbit anti-TH, Millipore Sigma, Cat#AB152, 1:750 in blocking buffer) at room temperature, later washed three times in PBS, and then incubated in secondary antibody (Alexa Fluor 488 Goat anti-Rabbit 1:1000, Cat#A11008) for 60 minutes. Tissue was then washed in 1x phosphate buffer (PB) and mounted on SuperFrost Plus microscope slides before being cover slipped with VECTASHIELD HardSet Antifade Mounting Medium with DAPI (Vector Laboratories, Cat# H-1500-10). Images were taken at 10x or 20x using a Nikon Ti2 epifluorescent microscope and fluorescence intensity of both the left and right striatum was quantified using QuPath software v0.4.3. Fluorescence intensity was quantified from three to five anatomically matched striatal sections per animal and averaged to generate a single value for each animal.

### Preparation of dissociated dorsal root ganglion neurons

Dorsal root ganglia (DRGs) were isolated using previously described procedures (44). Briefly, mice were deeply anesthetized with 5% isoflurane, and both left and right DRGs were collected and cleared of attached nerve roots and connective tissue. In 6-OHDA studies the DRGs were collected at least 10 days after intra-cranial administration of 6-OHDA to correspond with the pain-like behavior *in vivo*. Ganglia were enzymatically dissociated for 45 min at 37°C with gentle agitation in DMEM (Thermo Fisher Scientific, Cat#11965;) containing neutral protease (Worthington, Cat# LS02104, 1.04 mg/mL) and collagenase type I (Worthington, Cat#LS004194, 1.66 mg/mL;). Following digestion, tissue was mechanically dissociated by gentle trituration, and the resulting cell suspension was centrifuged to collect dissociated cells. Cells were resuspended in complete DRG culture medium consisting of DMEM supplemented with 10% fetal bovine serum (Fisher Scientific, Cat#FB12999105) and 1% penicillin/streptomycin (Life Technologies, Cat# 15140, 10,000 µg/mL stock). Dissociated cells were subsequently plated onto poly-D-lysine-coated glass coverslips and maintained in culture until electrophysiological recording.

### Measurement of Neuronal Excitability Using Whole-Cell Current-Clamp Electrophysiology

Electrophysiological recordings were conducted 16-24 h after plating using the whole-cell configuration of the patch-clamp technique in current-clamp mode. For experiments examining the effects of C194, dissociated DRG neurons were incubated with vehicle (DMSO) or C194 (5 μM) for 16-24 h before electrophysiological recording. Recordings were performed at RT (22–24 °C) using an EPC 10 amplifier (HEKA Elektronik; Germany) and acquired with PatchMaster v2x92 (HEKA). For current-clamp recordings, the external solution contained (in millimolar): 130 NaCl, 3 KCl, 2.5 CaCl_2_, 0.6 MgCl_2_, 10 D-Glucose, and 10 HEPES (pH 7.4 adjusted with NaOH, and mOsm/L = 300). The internal solution was composed of (in millimolar): 120 K-gluconate, 10 NaCl, 2 MgCl_2_, 5 EGTA, 4 Mg-ATP, and 10 HEPES (pH 7.3 mOsm/L = 300). DRG neurons were held at their resting membrane potential (RMP). Those neurons with RMP more hyperpolarized than -40 mV, stable baseline recordings, and evoked spikes that overshot 0 mV were used for experiments and analysis. Action potentials were evoked by applying a ramp from 0-250 pA over 1 s. Rheobase was measured from the ramp protocol by using the Easy Electrophysiology 2.7.3. software.

Pipettes were pulled from standard wall borosilicate glass capillaries (Sutter Instrument; Novato, CA, USA) with a horizontal puller (Model P-97, Sutter Instrument). The resistance of the pipettes when filled with internal solution and immersed in the recording bath ranged from 2 to 4 MΩ. Recordings were restricted to small-diameter DRG neurons with an average membrane capacitance of 13 pF (corresponding approximately to ∼20 μm diameter). Series resistance under 7 MΩ was deemed acceptable. Series resistance was compensated by 60–90%. Currents/signals were filtered at 10 kHz and digitized at 10-20 kHz. Analyses were performed using Fitmaster software v1.7 (HEKA) and Origin 9.0 software (OriginLab).

### Measurement of Voltage-Gated Sodium Currents Using Whole-Cell Voltage-Clamp Electrophysiology

Electrophysiological recordings were conducted 16-24 h after plating using the whole-cell patch-clamp technique in voltage-clamp mode. For experiments examining the effects of safinamide, dissociated DRG neurons were either exposed to safinamide (10 μM) or vehicle (0.1% DMSO) applied acutely in the external solution. Each coverslip was recorded up to 30 min. Recordings were performed at RT (22-24 °C) using an EPC 10 amplifier (HEKA Elektronik; Germany) and acquired with PatchMaster v2x92(HEKA). For voltage-clamp recordings of sodium currents, the external solution contained (in millimolar): 50 NaCl, 100 TEA-Cl, 1.8 CaCl_2_, 0.1 CdCl_2_, 1 MgCl_2_, 10 D-Glucose, and 10 HEPES (pH 7.3, and mOsm/L = 310); the reduced extracellular sodium concentration limited current amplitude to preserve voltage control, while TEA-Cl and CdCl_2_ blocked potassium and calcium currents, respectively. Glass pipettes were filled with the internal solution composed of (in millimolar): 140 CsF, 1.1 Cs-EGTA, 10 NaCl, and 10 HEPES (pH 7.3, and mOsm/L = 300). The capacitive transients were compensated, neurons were held at -90 mV, and only cells with adequate voltage control were used for experiments and analysis. Total Na_V_ currents were elicited by applying 150-millisecond voltage steps from -70 to +60 mV in 5-mV increments from a holding potential of -90 mV. For the groups that were treated with the Na_V_1.7 blocker ProTx-II, the toxin was present in the external solution at 5 nM. Steady-state inactivation (SSI) curves were obtained by applying 1-s conditioning pre-pulses from -120 to 10 mV in 10-mV increments followed by a 20-millisecond test pulse to -10 mV.

Sodium current densities were calculated by normalizing the current amplitude from each step with the cell capacitance. Inward currents were converted to conductance values using the equation

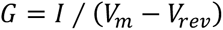

Where *G* is the conductance, *I* is the inward current, *V_m_* is the membrane potential step, *V_rev_* is the reversal potential determined for each cell.

Conductance data were normalized by the maximum conductance (*G_max_*) and fitted with the Boltzmann equation

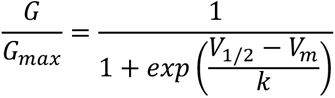

Where *V_1/2_* is the midpoint of activation and *k* is a slope factor.

Inactivation curves were obtained by dividing the peak current recorded at the test pulse by the maximum current (*I_max_*). Activation and SSI curves were fitted with the Boltzmann equation of the form.

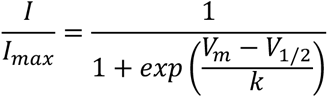

Where *V_m_* represents the inactivating pre-pulse membrane potential and *V_1/2_* represents the midpoint of fast inactivation.

### Molecular Modeling

Molecular docking of safinamide was performed against five human Na_V_1.7 cryo-EM structures (PDB: 7XM9, 7XMF, 7XMG, 8THG, 8THH) (45, 46) using Glide Standard Precision (Schrödinger Suite 2026-3) (47). Receptors were prepared with the Protein Preparation Wizard and ligands with LigPrep/Epik (48); receptor grids were centered on each structure’s co-crystallized ligand. For each receptor, the native ligand was redocked into its own site to benchmark pose accuracy (heavy-atom RMSD to the deposited coordinates), and safinamide was docked independently into the same grid. Full preparation, docking, and validation parameters are provided in Supplementary Methods. Figures were prepared using PyMOL (Schrödinger LLC, 2015).

### Human dorsal root ganglion preparation

All human tissue procurement procedures and ethical regulations were approved by the Institutional Review Board at the University of Texas at Dallas under protocol Legacy-MR-15-237. Fresh frozen human DRG samples were obtained from the Netherlands Brain Bank (NBB), Netherlands Institute for Neuroscience, Amsterdam (open access: www.brainbank.nl) and from organ donors through a collaboration with the Southwest Transplant Alliance (STA). All material was collected from donors who gave written informed consent and the use of the material and clinical information for research purposes was obtained by the NBB and by the STA. NBB donors typically enroll in the brain-donation program during life, enabling the collection of clinical information and post-mortem neuropathological assessment of conditions such as Parkinson’s disease. The STA obtains medical information on organ donors from recent hospital records and family-reported information. Demographic and diagnostic information for each donor are provided in **Supplementary Table S2**.

Fresh frozen DRGs were shipped on dry ice from the NBB to the University of Texas at Dallas and upon receipt, the samples were stored at -80°C. Upper thoracic DRGs were selected for use due to availability within the biobanks and to maintain level consistency between donors. All DRGs were embedded in OCT (Fisher Scientific, Cat#23-730-571) by applying small volumes within metal cryomolds over dry ice, allowing the medium to freeze incrementally so that the tissue never thawed. Cryoembedded samples were then sectioned on a cryostat. While in the cryostat chamber, the OCT was gently removed from the perimeter of each section with a paintbrush, and then the sections were placed into pre-chilled epitubes over dry ice. 100-μm worth of sections were placed into each tube to be used for RNA purification and sequencing. Residual OCT-blocks were wrapped in aluminum foil and returned to the -80°C freezer for later use.

### RNA purification and sequencing

Bulk RNA sequencing was used to identify transcriptomic differences in DRG associated with the Parkinson’s phenotype. For each ganglion, 100 μm sections were homogenized in QIAzol (Qiagen, Cat#79306) and RNA was recovered by chloroform phase separation using the RNeasy Plus Universal Mini Kit (Qiagen, Cat#73404). RNA concentration and purity were determined by NanoDrop and Qubit, and RNA integrity was measured on TapeStation, yielding RNA integrity numbers between 5.1-9.4. Libraries were prepared for paired-end mRNA sequencing and run on a NovaSeqX instrument at Psomagen as previously described (49). All donors passed quality control and were retained for downstream analysis.

### Human dorsal root ganglion RNA sequencing analysis

RNA-sequencing expression values from human thoracic DRG were quantified as transcripts per million (TPM) for individuals with PD (n = 8), age-matched controls (n = 8), and young controls (n = 4). TPM values were transformed as log2(TPM + 1) prior to statistical analysis. Genes were retained for each comparison if TPM was ≥1 in at least four samples within the relevant cohort. Principal component analysis was performed on the 2,000 most variable genes across all 20 donors using centered and scaled log2(TPM + 1) expression values. Differential expression between PD and age-matched controls was assessed using the limma package in R, with age included as a covariate in the linear model; sex was not included in the primary PD versus age-matched control model because both groups contained equal numbers of males and females. Empirical Bayes moderation was applied using eBayes, and model coefficients, moderated t statistics, nominal P values, and Benjamini-Hochberg false discovery rate-adjusted P values were calculated for all retained genes. Volcano plots displayed the limma model coefficient on the x axis and -log10(P) on the y axis, with genes meeting nominal P < 0.05 and an absolute effect-size threshold of log2(1.5) highlighted. The 30 genes with the smallest nominal P values in the PD versus age-matched control comparison were visualized by heatmap after row-wise Z-score transformation of log2(TPM + 1) expression values. In addition to the genome-wide analysis, expression of *DPYSL2* and *SCN9A* was examined based on the Na_V_1.7-CRMP2 mechanism investigated experimentally in the mouse studies. Expression of the HCN channel family (*HCN1*-*HCN4*) was additionally examined based on evidence that HCN-mediated current regulates Na_V_1.7 availability and Na_V_1.7-dependent DRG neuron excitability. Untransformed TPM values for each donor were plotted individually, with age-matched controls shown as open circles and PD samples as filled red circles; horizontal lines denote group means and error bars represent SEM. Exact P values displayed above gene-expression plots were derived from the age-adjusted limma model rather than from separate univariate tests. Additional three-group heatmaps were generated for curated ion-channel, neurotrophic/pain, immune/inflammatory, and nerve-injury/regeneration gene sets using row-wise Z scores, with samples displayed in the fixed order young controls, age-matched controls, and PD. All analyses and figures were generated in R using limma, ggplot2, ggrepel, dplyr, pheatmap, and openxlsx.

### Statistics

All data are expressed as mean ± standard error of the mean (SEM), with individual data points representing either single animals (behavioral and imaging experiments) or individual neurons (electrophysiological experiments). Statistical analyses were performed using GraphPad Prism (v10.2.3). As an additional quality-control step, electrophysiological datasets were screened using ROUT analysis (Q = 0.5%) prior to statistical analysis, with identified outliers excluded. Comparisons between two groups were performed using unpaired two-tailed t tests with Welch’s correction for unequal variances or Mann-Whitney U tests for nonparametric data, as appropriate. Comparisons involving more than two groups were analyzed using one-way ANOVA followed by either Tukey’s or Dunnett’s multiple comparisons tests. Factorial experiments were analyzed using two-way ANOVA, two-way repeated-measures ANOVA, three-way repeated-measures ANOVA, or mixed-effects models, as appropriate for the experimental design and presence of repeated measurements. When omnibus analyses revealed a significant interaction, appropriate post hoc multiple comparisons were performed using Tukey’s, Holm-Šidák’s, Šidák’s, Dunnett’s, or uncorrected Fisher’s least significant difference (LSD) tests, as appropriate. Statistical significance was defined as p < 0.05. No formal power analyses were performed. Sample sizes were determined based on prior experience with similar behavioral and electrophysiological assays.

### AI disclosure

Generative AI (ChatGPT, OpenAI; GPT-5 series, August-September 2026) was used to assist with development and refinement of R code for RNA-sequencing data analysis and with manuscript editing. All analyses, code, and resulting content were reviewed and verified by the authors. The authors assume full responsibility for the integrity and accuracy of AI-assisted output presented in this article.

### Graphics

Figures were generated using GraphPad Prism (v10.2.3), Adobe Illustrator 2022, and Biorender.com.

### Data Availability

Human DRG RNA-sequencing data generated in this study are being deposited in the NCBI Gene Expression Omnibus (GEO); accession information will be provided upon availability. Values underlying the graphical data presented in the manuscript and supplemental material are provided in the Supporting Data Values file. Additional data are available from the corresponding author upon reasonable request.

## Discussion

Pain is a common and disabling nonmotor symptom of Parkinson’s disease (PD), yet its underlying mechanisms remain poorly understood. Altered central nociceptive processing and loss of dopaminergic signaling have received the greatest attention, but dopamine replacement incompletely normalizes pain thresholds and sensory abnormalities are evident even in PD patients without chronic pain (50–53). Current models therefore implicate dysfunction across the pain neuraxis, including peripheral afferents, spinal and brainstem pathways, descending monoaminergic systems, basal ganglia, and cortical networks (10, 12). Here, we identify a peripheral component of this altered sensory state. Nigrostriatal neurodegeneration produced persistent hyperexcitability of primary sensory neurons associated with increased Na_V_1.7 current and altered sodium channel gating. Pharmacological disruption of CRMP2-dependent Na_V_1.7 regulation normalized sensory neuron excitability and reversed established pain, while genetic disruption of the Na_V_1.7 CRS prevented the development of pain for up to 30 weeks. These findings support peripheral sensory neuron sensitization as a mechanistic component of Parkinsonian pain and identify Na_V_1.7 and its regulation by CRMP2 as potential therapeutic targets.

Peripheral nervous system pathology is increasingly recognized in PD, but peripheral nerve degeneration and peripheral sensitization are not equivalent. Peripheral neuropathy has been identified in approximately 40% of well characterized PD cohorts, with small fiber abnormalities predominating (54). Reduced cutaneous and corneal sensory innervation is also present in early and levodopa naive PD, suggesting that peripheral pathology can accompany the disease itself rather than arising exclusively from treatment (55, 56). However, the relationship between fiber loss and pain is inconsistent. Peripheral abnormalities have been associated with hypersensitivity in some studies, whereas others report reduced sensory innervation together with increased thermal and mechanical thresholds (12, 56, 57). Thus, loss of sensory fibers alone does not adequately explain PD pain. Our findings identify a distinct peripheral process in which surviving primary sensory neurons become functionally hyperexcitable.

Zhang et al. provided important initial evidence for this possibility by demonstrating increased excitability and sodium current in small diameter DRG neurons following systemic MPTP administration (26). MPTP also increased *Scn9a* and *Scn10a* expression, while safinamide reduced DRG hyperexcitability and behavioral hypersensitivity (26). However, because MPTP was administered systemically, these experiments could not determine whether sensory neuron dysfunction occurred downstream of nigrostriatal neurodegeneration or resulted from direct or indirect peripheral consequences of the toxin. Our study addresses this question using 6-OHDA administered directly into the medial forebrain bundle, restricting the initiating neurotoxic insult to the CNS. Despite the absence of systemic neurotoxin exposure, small diameter DRG neurons developed increased sodium current and persistent hyperexcitability. To our knowledge, these findings provide the first evidence that neurodegeneration initiated within the central nigrostriatal system is sufficient to drive peripheral sensory neuron sensitization. This complements recent work showing that loss of the PD-associated gene *PARK7/DJ1* can act directly within sensory neurons to produce TRPA1-dependent hypersensitivity and peripheral neuropathy (58). Together, these findings suggest that peripheral sensory dysfunction in PD can arise both from intrinsic abnormalities within sensory neurons and as a secondary consequence of pathology initiated within the brain.

How a brain-restricted dopaminergic lesion produces peripheral sensitization remains unknown. Altered descending neural signaling, neuroendocrine responses, or systemic inflammatory signals accompanying nigrostriatal neurodegeneration could each contribute to this process. One possibility is that systemic responses accompanying neurodegeneration provide an early signal to peripheral nociceptors. Sensory neurons express receptors for cytokines and chemokines, and inflammatory signaling can regulate ion channel phosphorylation, trafficking, and neuronal excitability (59, 60). PD is associated with altered circulating inflammatory mediators and immune cell populations, providing a potential route for communication between central neurodegeneration and peripheral sensory neurons (57, 61). Importantly, central 6-OHDA lesions can themselves produce transient peripheral immune and neuroendocrine changes (62). An early systemic response to nigrostriatal degeneration could therefore prime nociceptor sensitization, whereas longer-lasting changes in ion channel function sustain hyperexcitability after that initiating signal resolves. This model is speculative but testable: early interventions should prevent sensitization, and channel-directed interventions should reverse it once established.

Our electrophysiological findings identify Na_V_1.7 as a major determinant of this persistent sensory state. ProTx-II eliminated the difference in sodium current between sham and 6-OHDA neurons, consistent with a substantial contribution of Na_V_1.7 to the increase in voltage-gated sodium conductance following 6-OHDA lesioning. In addition to increasing sodium current density, 6-OHDA altered the voltage dependence of sodium channel inactivation, indicating that nigrostriatal neurodegeneration produces functional remodeling of sodium currents beyond a simple increase in current magnitude. Na_V_1.7 is particularly important for nociceptor excitability because it amplifies subthreshold depolarizations and contributes to action potential initiation (27, 63). The increased Na_V_1.7-dependent current therefore provides a plausible mechanism for enhanced sensory neuron output following nigrostriatal neurodegeneration. Together, these findings suggest that Parkinsonian neurodegeneration alters peripheral sodium channel function through changes in current magnitude and biophysical regulation, although the mechanisms responsible for these changes remain unresolved.

Human genetics independently supports a relationship between Na_V_1.7 and PD pain. The nonsynonymous *SCN9A* rs6746030 variant has been associated with PD related pain, including central and musculoskeletal pain phenotypes (33). In an independent PD cohort, all five individuals carrying the heterozygous R1150W variant reported pain (34). R1150W produces an approximately 8-11 mV depolarizing shift in Na_V_1.7 activation and, when expressed in DRG neurons, depolarizes the resting membrane potential by approximately 6 mV and increases action potential firing approximately twofold in response to depolarization, providing a functional basis through which *SCN9A* variation could modify pain susceptibility (64, 65). More recently, severe refractory dysesthesias were reported in a patient with PD carrying an *SCN9A* variant, although this variant remains of uncertain significance (66). These observations do not establish Na_V_1.7 dysfunction as a universal mechanism of PD pain, but provide independent human evidence that Na_V_1.7 function can influence pain susceptibility in PD. Interestingly, *SCN9A* has also appeared in transcriptomic studies of other tissues in PD. *SCN9A* was among 29 differentially expressed genes identified in dermal fibroblasts from monozygotic twins discordant for PD, and increased *SCN9A* RNA was independently reported in extracellular RNA isolated from PD cerebrospinal fluid (67, 68). Although neither study examined pain or sensory neuron function, the independent identification of *SCN9A* in these datasets is notable in the context of the functional Na_V_1.7 phenotype identified here.

CRMP2-dependent regulation provides one mechanism through which Na_V_1.7 function can change independently of transcription. CRMP2 regulates Na_V_1.7 trafficking and membrane localization, and disruption of this pathway reduces Na_V_1.7 current and pathological pain (35–37, 69). The CRS within Na_V_1.7 is required for CRMP2-dependent regulation of the channel, and disruption of this domain reduces Na_V_1.7 current, membrane localization, presynaptic expression, and neuropathic pain while largely preserving physiological nociception (38). In the present study,

C194 normalized DRG hyperexcitability and reversed 6-OHDA-induced pain, while mice lacking the CRS were refractory to developing 6-OHDA-induced pain throughout the 30-week observation period. These complementary pharmacological and genetic findings support the Na_V_1.7 CRS as a critical determinant of the persistent PD pain-like phenotype and identify CRMP2-dependent Na_V_1.7 regulation as a therapeutically actionable mechanism. However, our data do not establish that nigrostriatal degeneration increases CRMP2 binding, CRMP2 SUMOylation, or Na_V_1.7 trafficking. Thus, although the CRS is required for the behavioral phenotype, the specific molecular event linking nigrostriatal degeneration to increased Na_V_1.7 function remains unresolved. Given the simultaneous changes in current magnitude and voltage-dependent inactivation, defining the underlying mechanism will require direct analysis of Na_V_1.7 membrane localization, channel biophysics, and posttranslational regulation.

Safinamide provides a clinically relevant link between Na_V_1.7 dysfunction and PD pain. Safinamide has multiple actions capable of influencing pain, including modulation of glutamatergic transmission and central monoaminergic signaling (19, 20), and our findings do not imply that peripheral Na_V_1.7 inhibition exclusively accounts for its analgesic efficacy. Multiple clinical studies have reported improvements in pain-related outcomes in PD patients by safinamide (14–18). Safinamide reduced concomitant analgesic use in pooled clinical analyses, and most improvement in pain-related quality of life was independent of reductions in OFF time, defined as periods of reduced dopaminergic treatment efficacy with re-emergence of Parkinsonian symptoms, as well as changes in depression or motor complications (14). Zhang et al. subsequently showed that safinamide directly suppresses DRG hyperexcitability induced by MPTP exposure (26). Our findings extend this work by identifying a ProTx-II sensitive Na_V_1.7 component of safinamide’s action in isolated sensory neurons. Because this effect occurs directly in isolated DRG neurons, it is independent of restoration of central dopaminergic signaling. These findings provide a potential peripheral mechanism for the reported analgesic actions of safinamide and further suggest that its therapeutic effects on pain extend beyond MAO-B inhibition. However, the contribution of peripheral Na_V_1.7 inhibition to analgesia in patients remains unknown.

The human DRG data provide translational context for the functional phenotype identified in mice. Consistent with the substantial overlap observed by PCA and the exploratory genome-wide analysis, PD DRG exhibited relatively limited global transcriptional remodeling, arguing against a broad transcriptional injury response as the primary explanation for the peripheral phenotype identified here.

Importantly, unchanged *SCN9A* and *DPYSL2* expression does not conflict with the functional mechanism identified in mice in our work. To our knowledge, neither *SCN9A* nor *DPYSL2* has emerged as a pain-associated differentially expressed transcript in the major human DRG RNA sequencing studies of painful diabetic neuropathy, neuropathic pain, or rheumatoid arthritis despite extensive transcriptional remodeling in these conditions (49, 70, 71). Indeed, *SCN9A* was robustly detected by Ray et al. and was used as a marker of neuronal RNA content, indicating that its absence from the pain-associated signature was not simply a detection problem (49). Human pain transcriptomics therefore provides little reason to expect increased Na_V_1.7 function to require increased *SCN9A* RNA. This is consistent with human *SCN9A* channelopathies, in which profound changes in pain arise from altered channel function, and with CRMP2-dependent regulation of Na_V_1.7 trafficking and membrane availability (27, 37, 72). Our human data therefore argue against transcriptional upregulation of *SCN9A* or *DPYSL2* in PD DRG, but do not test the functional properties highlighted by the mouse experiments, including channel gating, trafficking, membrane abundance, or posttranslational regulation. The mouse and human findings are therefore not discordant. Rather, together they point toward functional regulation of Na_V_1.7 as a mechanism that would not necessarily be captured by transcript abundance.

The HCN findings are particularly interesting in this context. *HCN2* and *HCN4* showed higher expression in human PD DRG, with *HCN3* showing a similar pattern, whereas *HCN1* was unchanged. HCN channels, particularly *HCN2*, have been implicated in inflammatory and neuropathic pain and can support repetitive nociceptor firing under some conditions (73). However, HCN currents do not uniformly increase sensory neuron excitability. Vasylyev et al. demonstrated that HCN current can increase rheobase, suppress subthreshold membrane potential oscillations, reduce repetitive firing, and counteract hyperexcitability produced by gain-of-function Na_V_1.7 (43). HCN current also altered Na_V_1.7 availability, providing a direct electrophysiological interaction between these conductances (43). The higher expression of multiple HCN family members in PD DRG is therefore unlikely to have a simple interpretation as a pro-excitatory pain signature. It could represent a contributor to altered sensory processing or a compensatory response that restrains sodium channel-dependent hyperexcitability. The latter possibility is particularly interesting because increased HCN conductance could oppose the increased Na_V_1.7-dependent excitability identified in our mouse model. However, transcript abundance cannot establish HCN current density or its net physiological effect, and direct electrophysiological studies of human PD sensory neurons will be required to distinguish between these possibilities.

Several limitations define the scope of these conclusions. The 6-OHDA model provides an important experimental advantage by isolating the consequences of a CNS dopaminergic lesion without systemic neurotoxin exposure, but it produces an acute nigrostriatal lesion and does not reproduce progressive α-synuclein pathology, multisystem neurodegeneration, or the prodromal phase of human PD (1, 74). Accordingly, these experiments address mechanisms capable of generating and maintaining sensory hypersensitivity following nigrostriatal neurodegeneration but do not model pain that can precede overt motor manifestations of PD. The behavioral assays used here measure stimulus-evoked sensory hypersensitivity and do not capture all dimensions of the spontaneous, musculoskeletal, dystonic, or central pain experienced by patients with PD. Because C194 is administered systemically and the ΔCRS manipulation is global, the behavioral experiments do not specifically localize the therapeutic or genetic effect to peripheral sensory neurons, even though the DRG data demonstrate a peripheral phenotype. We also do not identify the signal connecting nigrostriatal degeneration to NaV1.7 dysfunction or establish the molecular mechanism responsible for the altered NaV1.7 current and gating. Finally, human bulk RNA sequencing measures transcript abundance rather than ion channel physiology, and donor pain histories were unavailable. Accordingly, the human data establish molecular changes in PD DRG but cannot determine which changes are specifically associated with pain. Direct functional analysis of human PD sensory neurons and larger human cohorts with detailed pain phenotyping will be important next steps.

Together, these findings establish a mechanistic connection between central Parkinsonian neurodegeneration and peripheral sensory neuron dysfunction. Previous work has demonstrated peripheral fiber loss in PD, intrinsic sensory dysfunction in genetic PD models, DRG hyperexcitability following systemic MPTP, and associations between functional *SCN9A* variants and PD pain (26, 33, 34, 56, 58). Here we demonstrate that neurodegeneration initiated within the brain is sufficient to produce persistent Na_V_1.7-dependent hyperexcitability in primary sensory neurons and identify the CRMP2-Na_V_1.7 regulatory axis as a critical determinant of this pain state. The human data further suggest that peripheral sensory dysfunction in PD can occur without transcriptional induction of *SCN9A* or *DPYSL2* and raise the possibility that altered HCN channel expression modifies peripheral sensory excitability. Together, these findings expand the mechanistic framework for Parkinsonian pain beyond the CNS to include primary sensory neurons and identify Na_V_1.7 regulation as an actionable peripheral mechanism for therapeutic intervention.

## Acknowledgements

This work was supported by National Institutes of Health (NIH) awards K00NS124190 (to T.S.N.), F32NS128392 (to H.N.A.), K99NS134965 (to K.G.), T32NS082128 (to A.L.W.), and RF1NS131165, R61NS126026, and R01NS120663 (to R.K.); by an American Neuromuscular Foundation Development Grant (to T.S.N.); by a PhRMA Foundation Postdoctoral Fellowship in Drug Discovery 1335819 (to E.J.R.-P.); and by a Parkinson’s Foundation Summer Student Fellowship (to N.K.G.). Generation of the Na_V_1.7-CRS mouse line was made possible by a pilot grant from the Genetically Engineered Mouse Models (GEMM) Core at the University of Arizona (to R.K.). We thank Dr. Teodora G. Georgieva for generating the mouse line.

## Author contributions

TSN and RK conceived and designed the study. TSN, NKG, HNA, and EJRP performed behavioral experiments, surgeries, and imaging studies. ACR, SLL, ALW, and KG performed electrophysiological experiments. EE designed, performed, and analyzed the molecular docking studies. TSN, SIS, IS, and TJP contributed to the human DRG RNA-sequencing studies, with TJP overseeing the RNA-sequencing experiments. TSN performed the downstream RNA-sequencing analysis and generated the corresponding figures. TSN, NKG, EJRP, ALW, EE, and IS contributed to data analysis and integration. TSN prepared the figures and wrote the initial manuscript. TSN and RK wrote and edited the manuscript. RK supervised the overall study, with TSN contributing to supervision of experimental work. TSN, NKG, HNA, EJRP, ALW, KG, RK, and TJP acquired funding. All authors contributed to interpretation of the data, reviewed and edited the manuscript, and approved the final version. Relative contributions are graphically illustrated below.

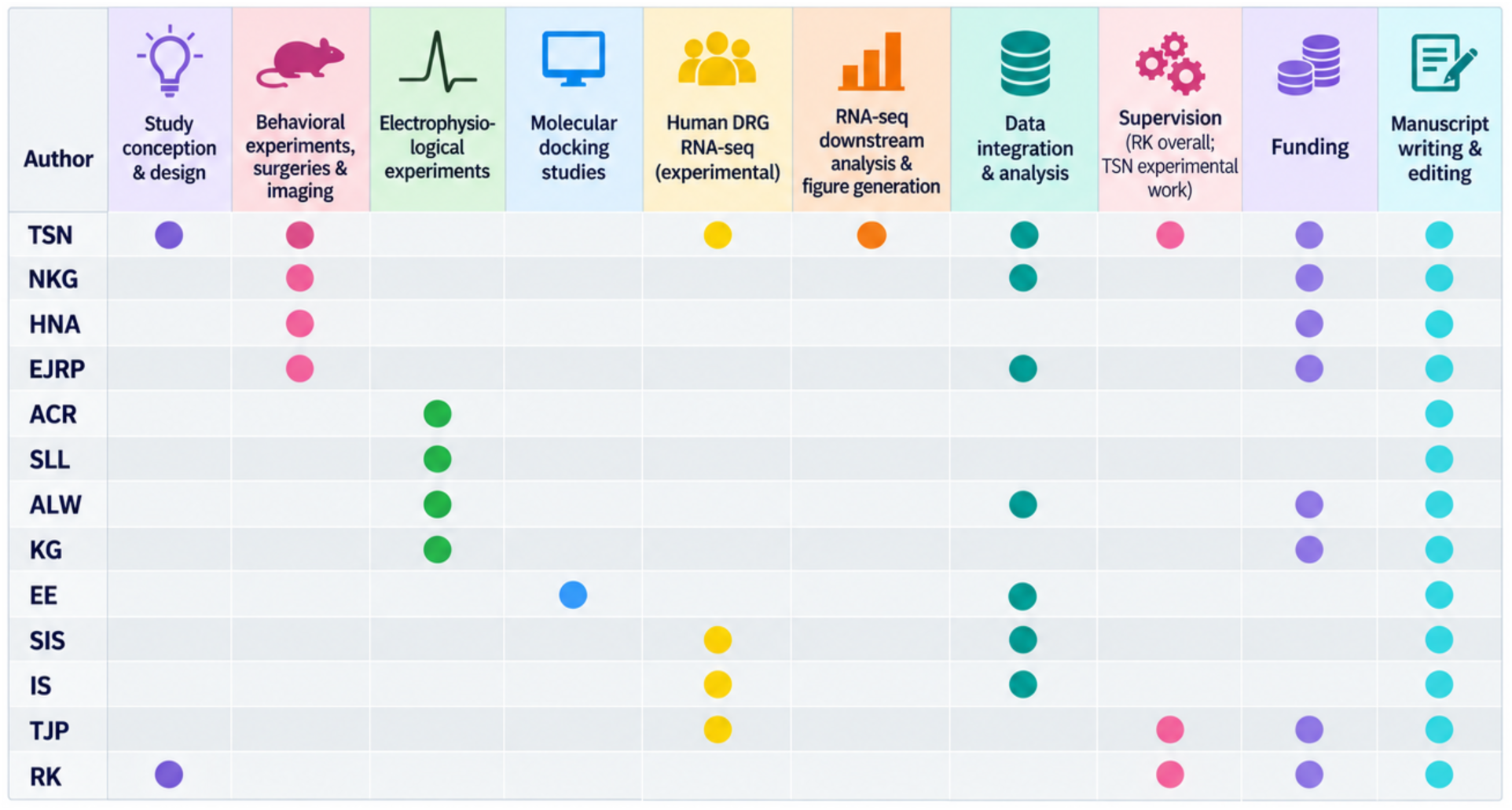

