## Supplementary Material for "Na_V_1.7-dependent peripheral sensitization drives chronic pain in Parkinson’s disease"

**Supplementary Methods**

**Receptor and Ligand preparation**

Five human $Na_V1.7$ cryo-EM structures: 7XM9, 7XMF, 7XMG, 8THG, and 8THH were imported into Maestro and processed with the Protein Preparation Wizard(1): bond orders were assigned, hydrogens were added, missing side chains within the binding region were checked for completeness, and het states (including the co-crystallized ligand) were generated with Epik at pH 7.4 ± 2.0. Hydrogen-bond assignment was optimized and protonation states calculated with PROPKA at pH 7.4(2). The structure was subjected to a restrained minimization with the OPLS4 force field(3, 4), converging heavy atoms to 0.30 Å RMSD. Water molecules were removed. Receptors were treated as rigid throughout docking.

(S)-Safinamide was constructed with the (S) configuration at its single stereocenter explicitly specified; the enantiomer was not enumerated. Ligands were processed with LigPrep(1), generating ionization states at pH 7.4 ± 2.0 with Epik(5), retaining specified chiralities, generating low-energy ring conformations, and using OPLS4 for energy minimization. Both neutral and protonated Epik-generated states of safinamide were docked to assess the effect of ionization on binding. Reference ligands (XEN907, $Na_V1.7$-IN2, TC-N1752, riluzole, lamotrigine) were extracted from their deposited coordinates and processed identically for redocking.

Receptor grids were generated in Maestro, centered on the centroid of each structure's co-crystallized ligand. The outer box was enlarged to 20 Å per side to accommodate safinamide's conformation, adjusting the ligand-length settings so extended poses were not truncated during sampling.

**Molecular Docking**

Docking was performed with Glide SP Schrödinger Release 2026-3(6). For each receptor, the extracted co-resolved ligand was redocked into its own grid, and safinamide was docked independently into the same grid. Flexible ligand sampling was used with Epik state penalties added to the docking score; up to 5 poses per ligand were retained, with post-docking minimization enabled. The top-ranked pose by Glide Score was retained per ligand receptor pair. Heavy-atom RMSD between each redocked reference-ligand pose and its deposited coordinates was computed using VMD(7).

### Supplementary Results

We used molecular docking to test whether safinamide is structurally compatible with a defined drug-binding site in $Na_V1.7$, and whether this compatibility is robust across different receptor conformations. Redocking of each structure's co-resolved ligand into its own receptor, using Glide SP, reproduced the deposited pose within the conventional heavy-atom RMSD threshold in three of five structures: 8THH (0.574 Å), 7XM9 (1.356 Å), and 8THG (1.428 Å). Two structures exceeded this threshold slightly: 7XMF (2.086 Å) and 7XMG (2.443 Å), both co-resolved with the largest and most conformationally flexible reference ligands in the set ($Na_V1.7$-IN2 and TC-N1752, respectively)(8). Safinamide's top-ranked docked pose occupied the central pore cavity in all five structures examined (**Fig. S1**), including the two structures with higher redocking RMSD, overlapping the region occupied by the co-resolved ligand in each case. Contact analysis identified the $D_{IV}$-S6 residue F1748 as the residue engaged by safinamide, via π-stacking, in every structure examined (**Fig. S1, C-G**). This residue corresponds to the aromatic residue of the canonical local-anesthetic receptor site conserved across voltage-gated sodium channels and is structurally homologous to the residue at which the p.F1586C substitution in $Na_V1.4$ was previously shown to substantially impair safinamide block(9), providing an independent, mutagenesis-grounded rationale for the pocket identified.

Safinamide's binding mode converges on a small set of conserved, high-confidence contacts, most notably a $D_{III}$ cluster (W1332, L1329, C1328) engaged consistently in the two best-validated structures with the most chemically comparable reference ligands (**Fig. S1**). Beyond the shared contact described above, safinamide formed hydrogen bonds with T1695/T1696 in four structures (7XMF, 7XMG, 8THG, 8THH) and with Q360 in two structures (7XMF, 8THH). It also formed a halogen-bond-consistent interaction between its meta-fluorophenyl substituent and this $D_{III}$ cluster in 7XMG and 8THG, with an analogous contact at a neighboring position in 7XM9. Because this cluster sits adjacent to the $D_{IV}$-S6 local-anesthetic residue and was reproducibly engaged in the two structures with the highest redocking performance (riluzole and lamotrigine, RMSD 1.428 Å and 0.574 Å, respectively) (**Table S1**), we consider these two complexes the most informative comparators for interpreting safinamide's binding mode, given their similar size, polarity, and validation quality. Safinamide is favored relative to the native ligands across most of the structures examined: it produced more favorable GlideScores than the corresponding resolved ligand in four of five receptor structures, with the largest differences seen for the two best-validated small-molecule reference complexes, including riluzole (**Table S1**). Because GlideScores are affected by ligand size, flexibility, and pocket complementarity, these results should be interpreted only as indicating that safinamide is not disfavored relative to each structure's native ligand, not as evidence of superior binding affinity. These calculations model a rigid receptor and do not incorporate side-chain flexibility beyond the docking search itself, membrane environment, explicit solvent, or entropic contributions to binding. The site identified here is highly conserved across $Na_V$ subtypes, and these results should not be interpreted as evidence of $Na_V1.7$ selectivity.

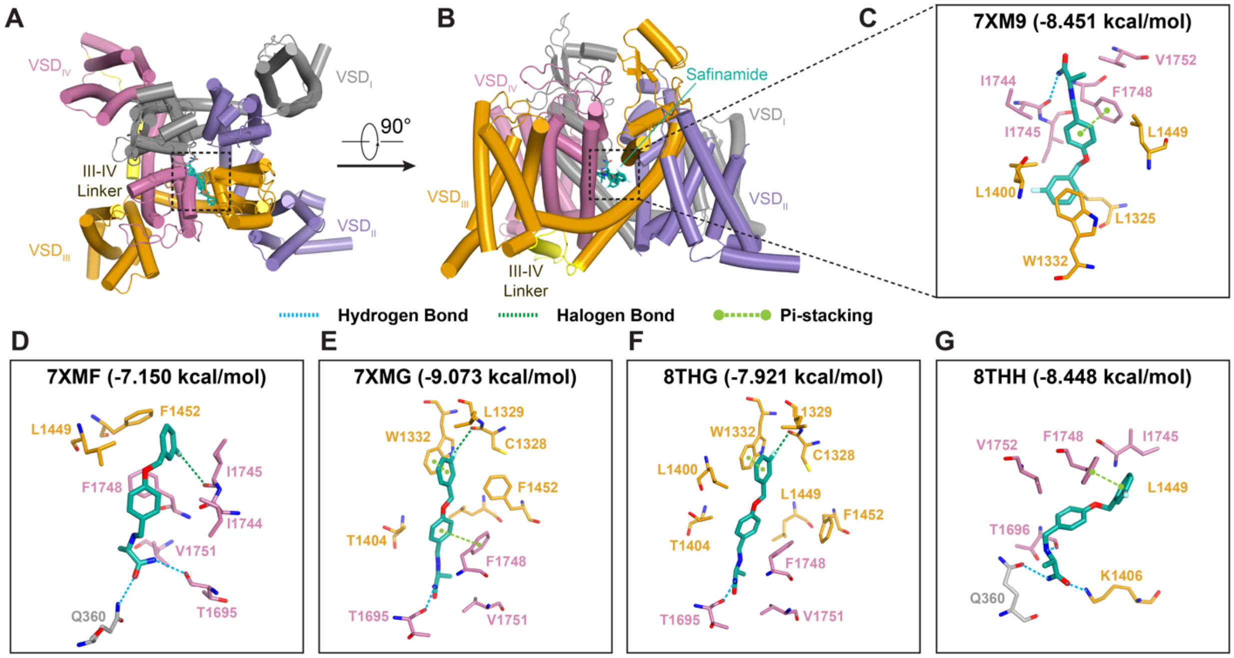


**Supplementary Figure S1. Molecular docking of safinamide within the Na$_V$1.7 central cavity, overlapping the binding site of sodium channel blockers. (A)** Extracellular (top-view) of the human Na$_V$1.7 (shown for PDB 7XM9), depicting the four repeats (D$_I$, grey; D$_{II}$, light purple; D$_{III}$, orange; D$_{IV}$, pink) with the D$_{III}$–D$_{IV}$ linker (yellow). Safinamide (teal licorice) is docked within the central cavity formed by the S6 helices of the four repeats; dashed box indicates the region magnified in panels C-G. **(B)** The same complex viewed after a 90° rotation about the membrane-normal axis, oriented with the extracellular side up; the safinamide docking pose is labeled for clarity. **(C-G)** Close-up views of the top-ranked safinamide binding pose and contacting residues in each of five Na$_V$1.7 structures. Docking was performed in Maestro using Glide; the GlideScore of the top-ranked pose (kcal/mol) is shown above each panel. Interaction type is indicated by dashed lines: hydrogen bond (blue), halogen bond (dark green), π-stacking (green, terminal circles).

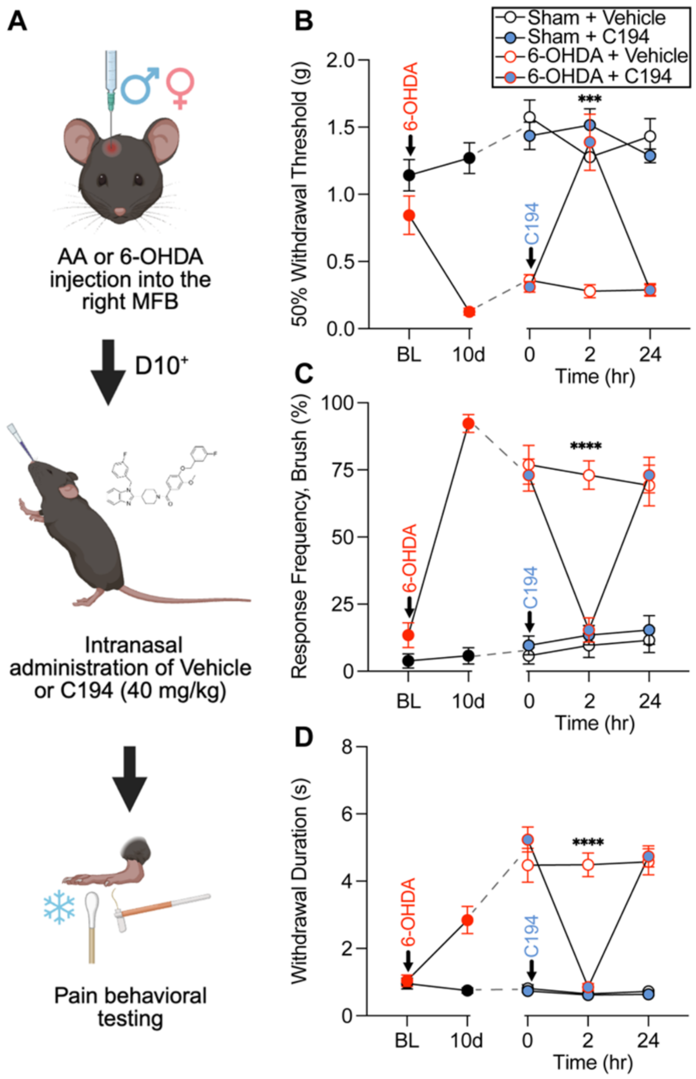


**Supplementary Figure S2. Intranasal administration of Compound 194 reverses established Parkinsonian pain-like behavior. (A)** Experimental design for intranasal Compound 194 (C194) administration. Beginning ≥10 days following unilateral injection of ascorbic acid (AA; sham) or 6-hydroxydopamine (6-OHDA) into the right medial forebrain bundle (MFB), male and female mice received a single 20 µl intranasal administration of vehicle (DMSO) or C194 (40 mg/kg), followed by pain behavioral testing over 24 h. **(B)** Mechanical withdrawal thresholds measured using von Frey (vF) filaments, **(C)** brush-evoked response frequency, and **(D)** acetone-evoked withdrawal duration in sham and 6-OHDA mice following intranasal vehicle or C194 administration (n = 13 mice/group). Intranasal C194 reversed 6-OHDA-induced mechanical hypersensitivity, dynamic mechanical allodynia, and cold hypersensitivity. For **B–D**, statistical comparisons were performed using two-way repeated measures ANOVA followed by Tukey's multiple-comparisons test. Data are presented as mean ± SEM. *** $P < 0.001$, **** $P < 0.0001$.

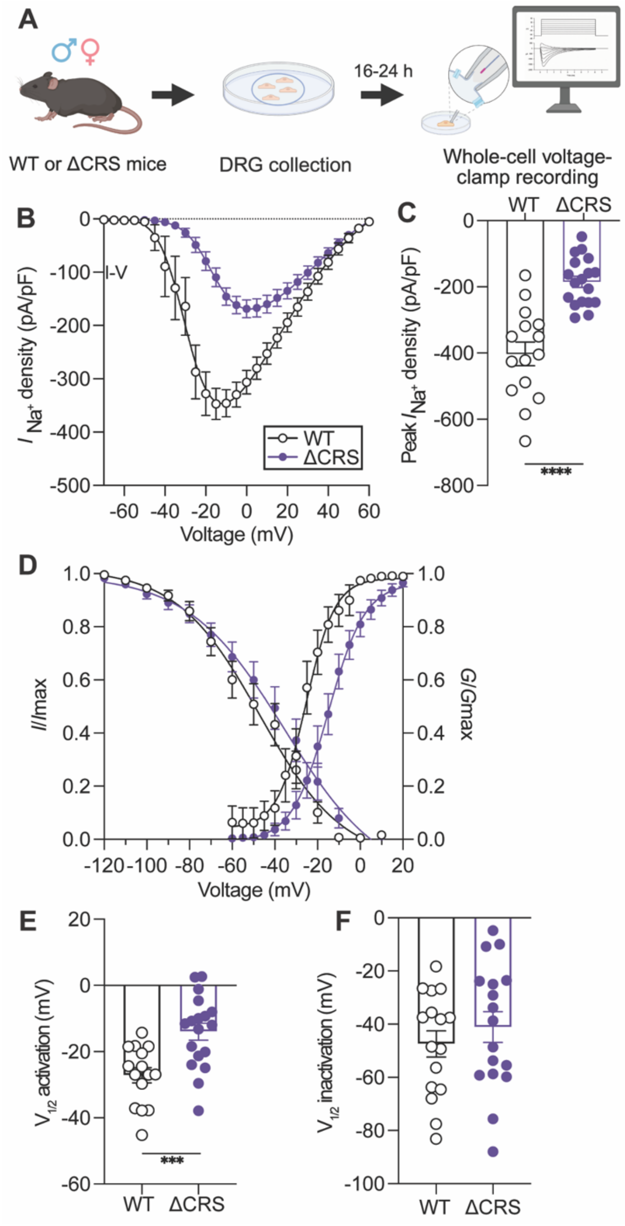


**Supplementary Figure S3. Disruption of the $Na_V1.7$ CRMP2 regulatory sequence reduces sodium current density and alters voltage-dependent gating in primary sensory neurons.** (**A**) Experimental design for whole-cell voltage-clamp recordings from dorsal root ganglion (DRG) neurons isolated from naïve wild-type (WT) and $Na_V1.7^{CRS\Delta/\Delta}$ (ΔCRS) mice. DRG neurons were cultured for 16-24 h prior to recording. (**B**) Current-voltage (I-V) relationships for total voltage-gated sodium current in small-diameter DRG neurons from WT and ΔCRS mice. (**C**) Peak sodium current density was significantly reduced in ΔCRS neurons compared with WT neurons. (**D**) Voltage dependence of sodium channel activation and steady-state inactivation in WT and ΔCRS neurons. (**E**) Half-maximal voltage of activation ($V_{1/2}$ activation) was significantly shifted in ΔCRS neurons compared with WT neurons. (**F**) Half-maximal voltage of steady-state inactivation ($V_{1/2}$ inactivation) in WT and ΔCRS neurons. B, E, and F were analyzed with an unpaired two tailed t test with Welch’s correction. n = 15-18 cells/group. Data are presented as mean ± SEM with individual cells shown where applicable. $^{**}P < 0.01$, $^{***}P < 0.001$.

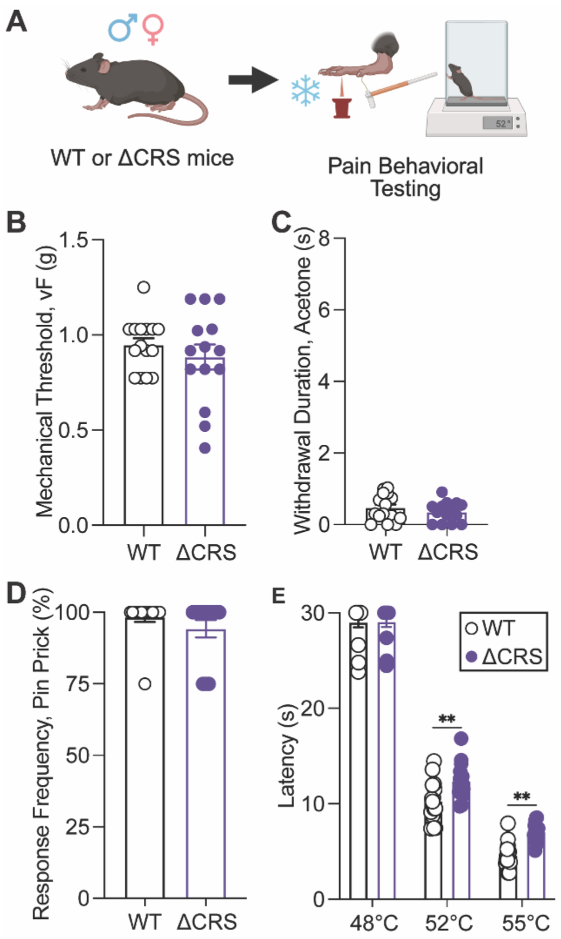


**Supplementary Figure S4. Baseline somatosensory function is largely preserved following disruption of the Na$_V$1.7 CRMP2 regulatory sequence.** (**A**) Experimental design for assessment of baseline somatosensory function in naïve WT and ΔCRS mice. (**B**) Mechanical withdrawal thresholds assessed using von Frey testing. (**C**) Cold sensitivity assessed by acetone-evoked withdrawal duration. (**D**) Response frequency to a blunt pin prick. (**E**) Thermal withdrawal latencies assessed using hot-plate testing at 48°C, 52°C, and 55°C. ΔCRS mice exhibited largely preserved baseline somatosensory function, with modest differences in thermal sensitivity at higher temperatures. **B-D** were analyzed using unpaired two-tailed *t* tests. **E** was analyzed using two-way repeated-measures ANOVA followed by Šidák's multiple comparisons test. n = 14-15 mice/group. Data are presented as mean ± SEM with individual animals shown. **P < 0.01

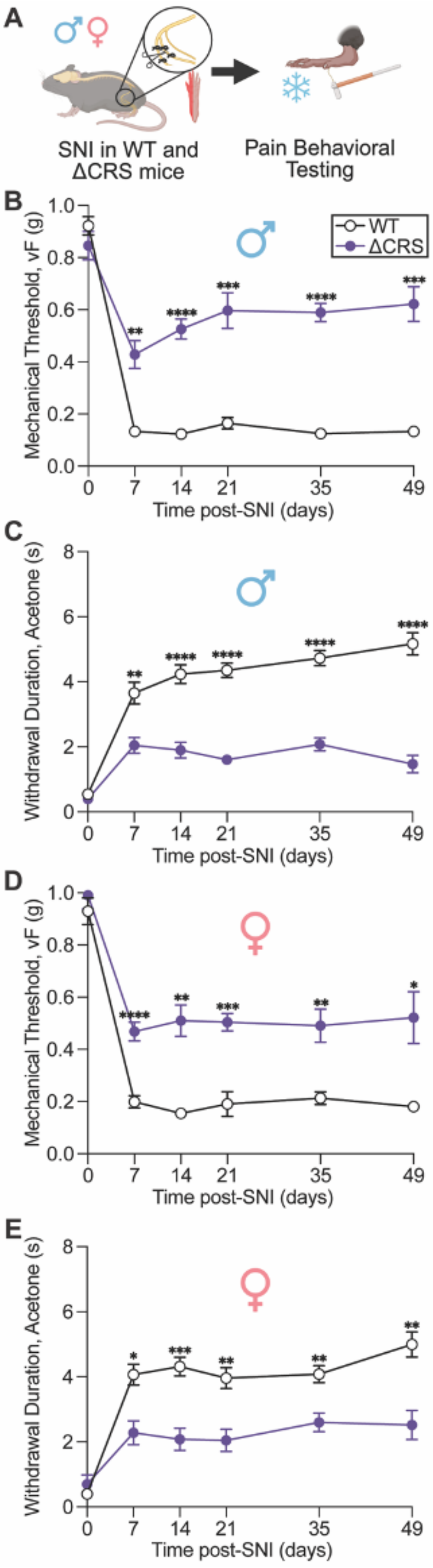
A
SNI in WT and ΔCRS mice
Pain Behavioral Testing
B
Mechanical Threshold, vF (g)
WT
ΔCRS
Time post-SNI (days)
C
Withdrawal Duration, Acetone (s)
Time post-SNI (days)
D
Mechanical Threshold, vF (g)
Time post-SNI (days)
E
Withdrawal Duration, Acetone (s)
Time post-SNI (days)

**Supplementary Figure S5. Genetic disruption of the $Na_V1.7$ CRMP2 regulatory sequence attenuates neuropathic pain-like behavior following spared nerve injury**. (**A**) Experimental design for spared nerve injury (SNI) and longitudinal assessment of mechanical and cold sensitivity in male and female WT and ΔCRS mice. (**B**) Mechanical withdrawal thresholds following SNI in male WT and ΔCRS mice. (**C**) Acetone-evoked withdrawal duration following SNI in male WT and ΔCRS mice. (**D**) Mechanical withdrawal thresholds following SNI in female WT and ΔCRS mice. (**E**) Acetone-evoked withdrawal duration following SNI in female WT and ΔCRS mice. ΔCRS mice exhibited markedly attenuated mechanical and cold hypersensitivity following SNI in both sexes compared with WT mice. **B-E** were analyzed using two-way repeated-measures ANOVA followed by Šidák's multiple comparisons test. n = 10 mice/group. Data are presented as mean ± SEM. *P < 0.05, **P < 0.01, ***P < 0.001, ****P < 0.0001.

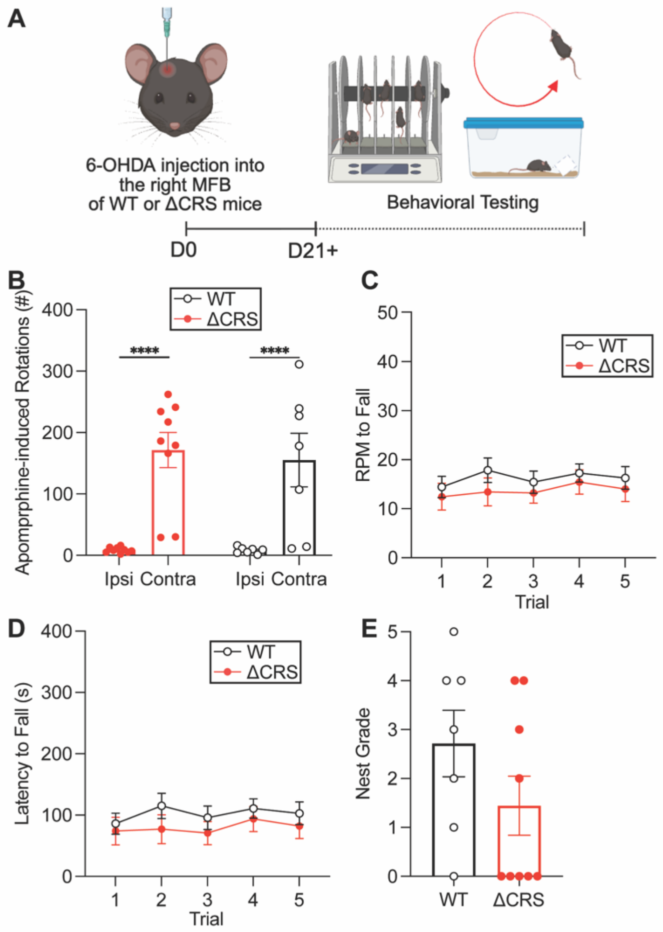


**Supplementary Figure S6. Na$_V$1.7 CRS disruption does not alter motor deficits following nigrostriatal dopaminergic neurodegeneration. (A)** Experimental design for motor behavioral assessment following unilateral injection of 6-hydroxydopamine (6-OHDA) into the right medial forebrain bundle (MFB) of wild-type (WT) and Na$_V$1.7$^{CRS\Delta/\Delta}$ (ΔCRS) mice. Behavioral testing was performed beginning ≥21 days following 6-OHDA lesion. **(B)** Apomorphine-induced ipsilateral and contralateral rotations in WT and ΔCRS mice. **(C)** Rotarod performance expressed as revolutions per minute (RPM) at fall across five trials. **(D)** Rotarod latency to fall across five trials. **(E)** Nest-building performance in WT and ΔCRS mice. No significant differences in apomorphine-induced rotations, rotarod performance, or nest-building behavior were observed between WT and ΔCRS mice. Data in **B** was analyzed with a two-way ANOVA followed by Uncorrected Fisher's LSD multiple comparisons test, **E** was analyzed using unpaired two-tailed *t* tests, and data in **C** and **D** were analyzed using two-way repeated-measures ANOVA followed by Šidák's multiple comparisons test. n = 7-9 mice/group. Data are presented as mean ± SEM with individual animals shown where applicable.

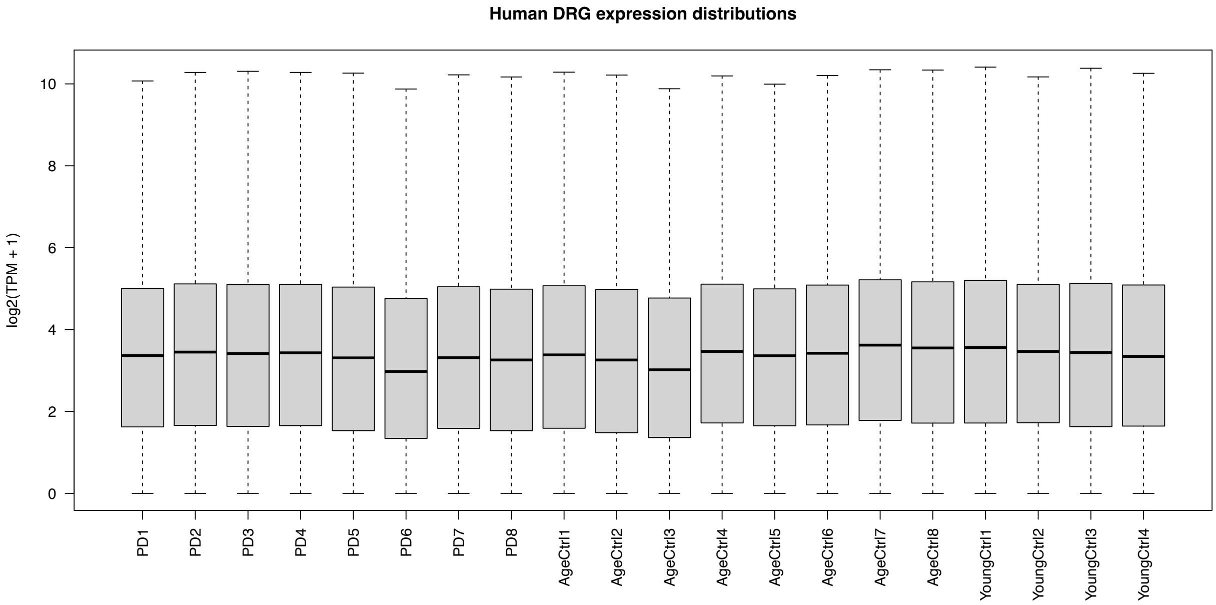


**Supplementary Figure S7. Gene expression distributions across human dorsal root ganglion RNA-sequencing samples.** Boxplots showing the distribution of log2(TPM + 1) gene expression values for each human dorsal root ganglion (DRG) sample included in the transcriptomic analysis, comprising Parkinson's disease (PD; n = 8), age-matched control (n = 8), and young control (n = 4) donors. Similar expression distributions were observed across samples and diagnostic groups. Boxes represent the interquartile range with the median indicated by the center line; whiskers represent the distribution of expression values.

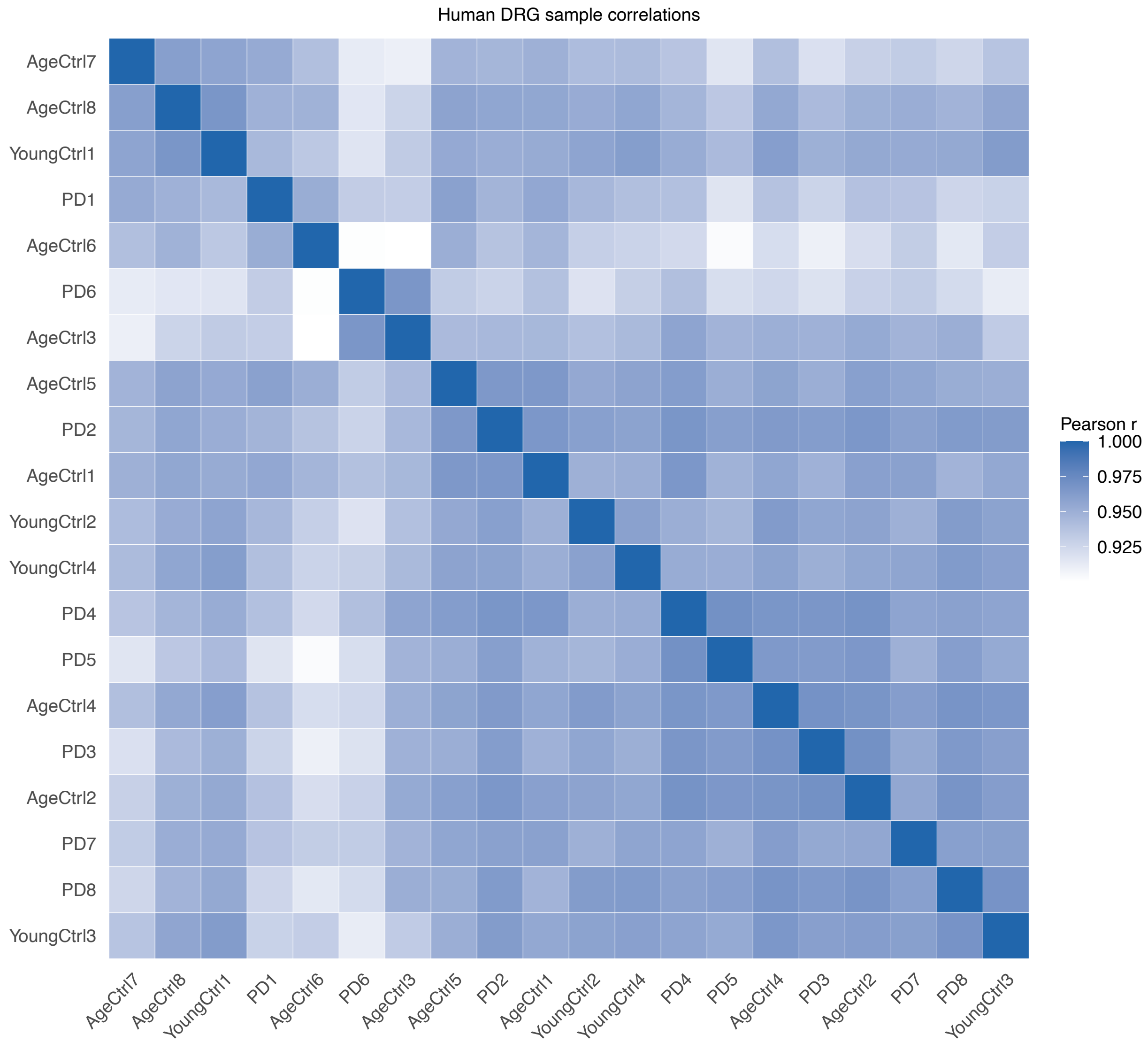


**Supplementary Figure S8. Correlation of gene expression profiles across human dorsal root ganglion RNA-sequencing samples.** Heatmap showing pairwise Pearson correlation coefficients among human dorsal root ganglion (DRG) transcriptomes from individuals with Parkinson's disease (PD; n = 8), age-matched controls (n = 8), and young controls (n = 4). Pearson correlations were calculated using log2(TPM + 1) expression values across genes. High correlations were observed across all samples, with no individual sample exhibiting a markedly divergent global expression profile. Samples are ordered by hierarchical clustering.

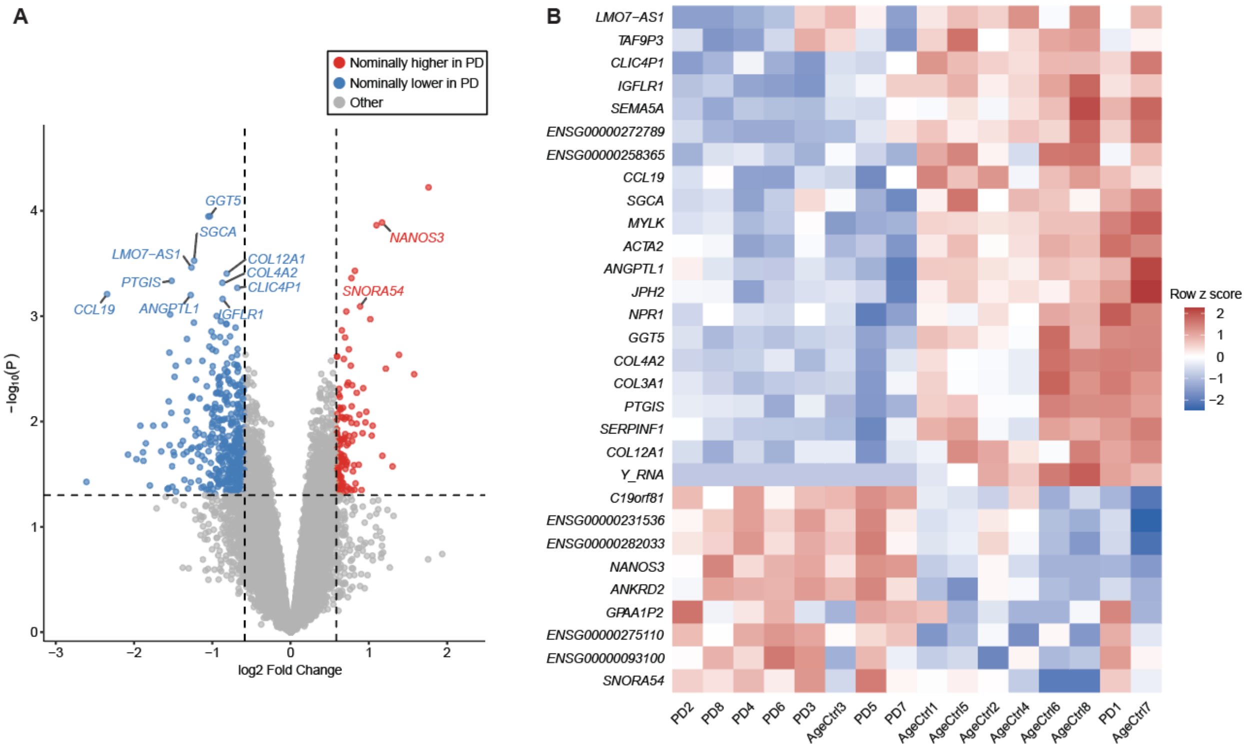


**Supplementary Figure S9. Exploratory genome-wide transcriptional analysis of human Parkinson's disease dorsal root ganglia. (A)** Volcano plot of differential gene expression between Parkinson's disease (PD; n = 8) and age-matched control (n = 8) dorsal root ganglia (DRG). Differential expression was assessed using limma with age included as a covariate and empirical Bayes moderation. The x axis represents the limma model coefficient and the y axis represents −log10(*P*). Genes meeting nominal $P < 0.05$ and an absolute effect-size threshold of log2(1.5) are highlighted. **(B)** Heatmap showing the 30 genes with the smallest nominal *P* values from the PD versus age-matched control comparison. Expression values are displayed as row-wise Z scores of log2(TPM + 1), with individual donors shown as columns and genes shown as rows.

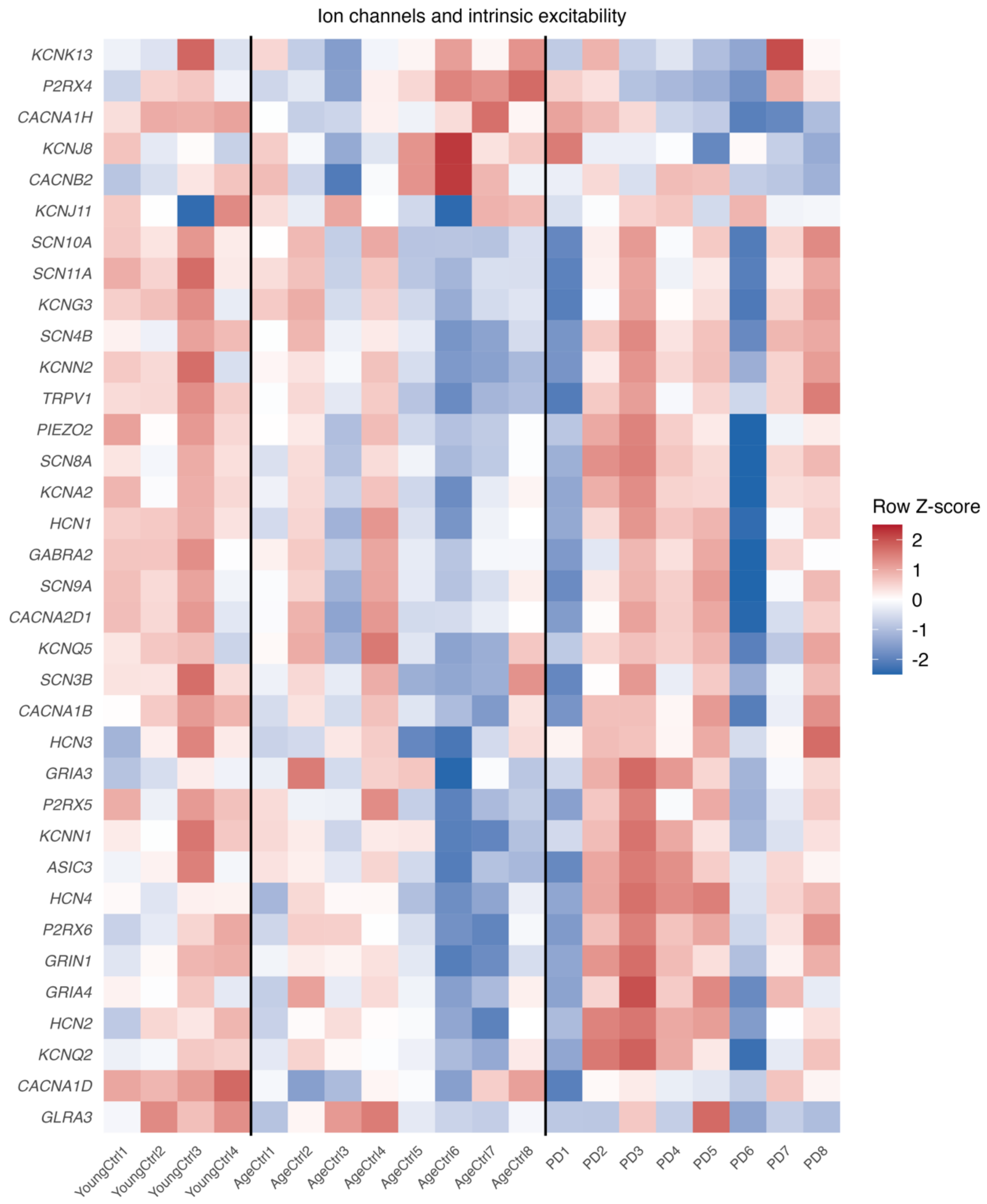


**Supplementary Figure S10. Expression patterns of ion channel and intrinsic excitability genes across young control, age-matched control, and Parkinson's disease human dorsal root ganglia.** Heatmap showing expression of candidate genes associated with ion channel function and intrinsic neuronal excitability in human dorsal root ganglia (DRG) from young controls (n = 4), age-matched controls (n = 8), and individuals with Parkinson's disease (PD; n = 8). Expression values were log2(TPM + 1) transformed and standardized by row across donors to generate Z scores, with higher and lower relative expression indicated for each gene. Samples are grouped by donor cohort, with vertical lines separating young controls, age-matched controls, and PD donors. The heatmap provides a descriptive comparison of expression patterns across age and disease groups; statistical comparisons between PD and age-matched controls were performed separately using the age-adjusted limma model.

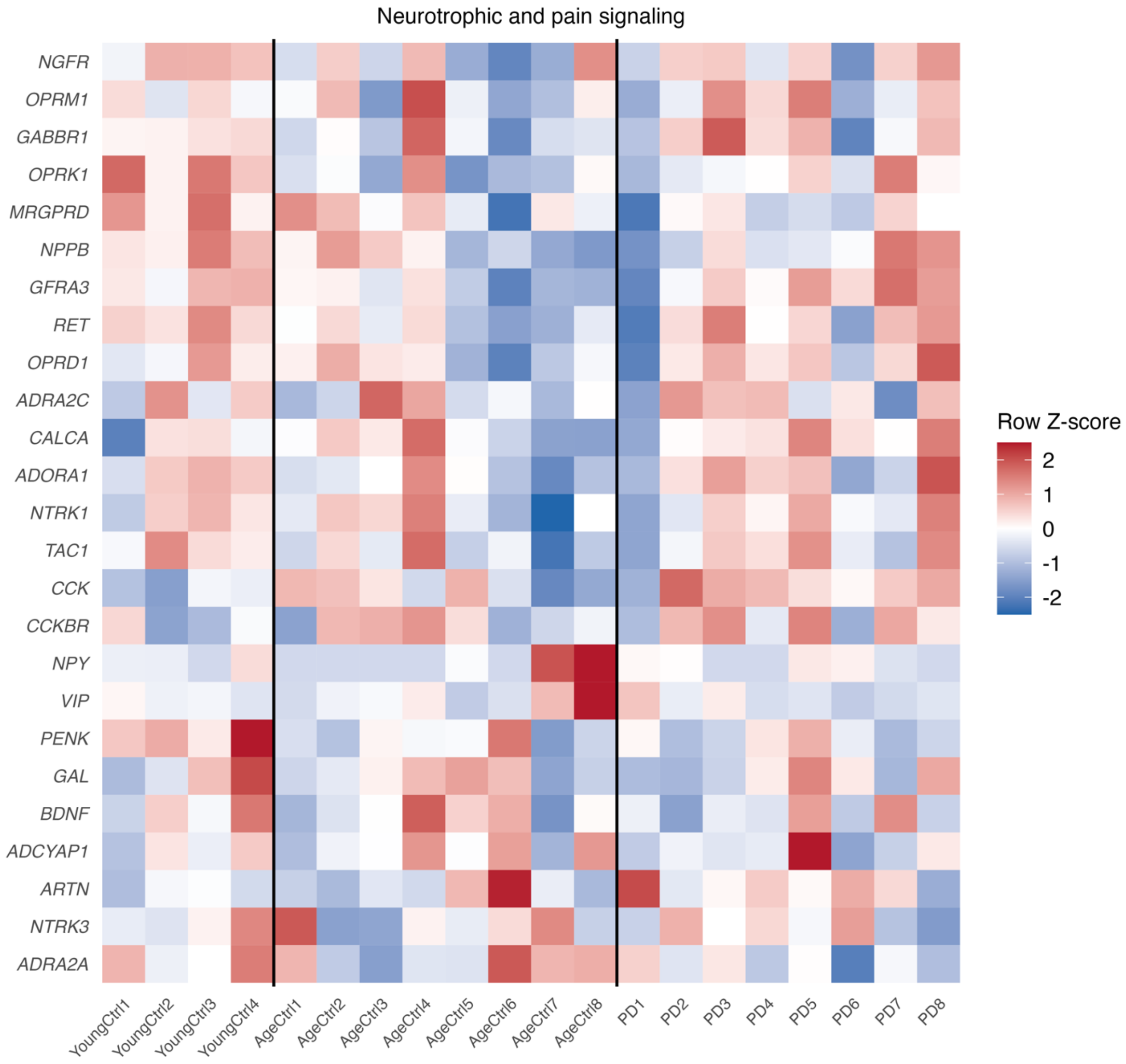


**Supplementary Figure S11. Expression patterns of neurotrophic and pain signaling genes across young control, age-matched control, and Parkinson's disease human dorsal root ganglia.** Heatmap showing expression of candidate genes associated with neurotrophic and pain signaling in human dorsal root ganglia (DRG) from young controls (n = 4), age-matched controls (n = 8), and individuals with Parkinson's disease (PD; n = 8). Expression values were log2(TPM + 1) transformed and standardized by row across donors to generate Z scores, with higher and lower relative expression indicated for each gene. Samples are grouped by donor cohort, with vertical lines separating young controls, age-matched controls, and PD donors. The heatmap provides a descriptive comparison of expression patterns across age and disease groups; statistical comparisons between PD and age-matched controls were performed separately using the age-adjusted limma model.

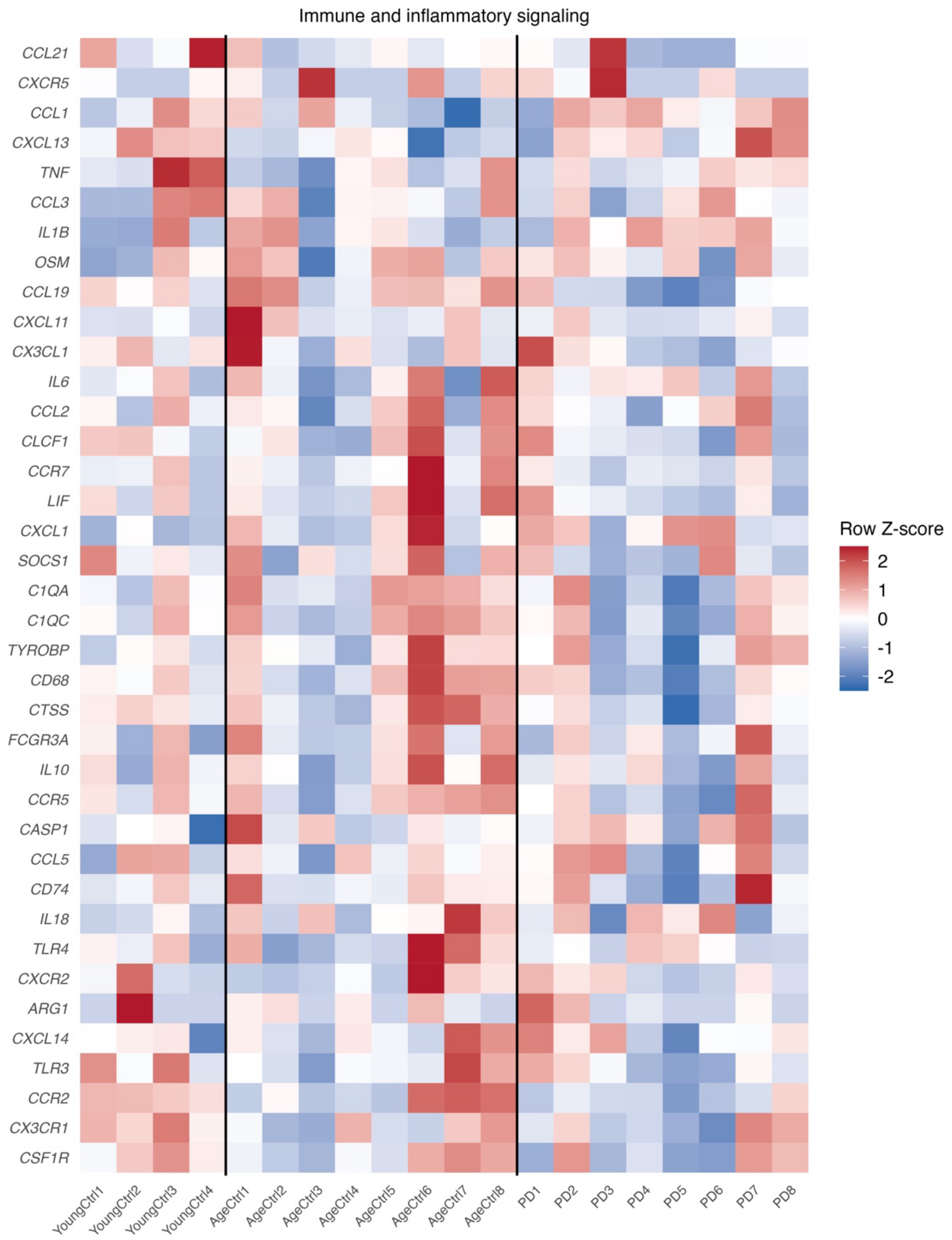


**Supplementary Figure S12. Expression patterns of immune and inflammatory signaling genes across young control, age-matched control, and Parkinson's disease human dorsal root ganglia.** Heatmap showing expression of candidate genes associated with immune and inflammatory signaling in human dorsal root ganglia (DRG) from young controls (n = 4), age-matched controls (n = 8), and individuals with Parkinson's disease (PD; n = 8). Expression values were log2(TPM + 1) transformed and standardized by row across donors to generate Z scores, with higher and lower relative expression indicated for each gene. Samples are grouped by donor cohort, with vertical lines separating young controls, age-matched controls, and PD donors. The heatmap provides a descriptive comparison of inflammatory and immune-related expression patterns across age and disease groups; statistical comparisons between PD and age-matched controls were performed separately using the age-adjusted limma model.

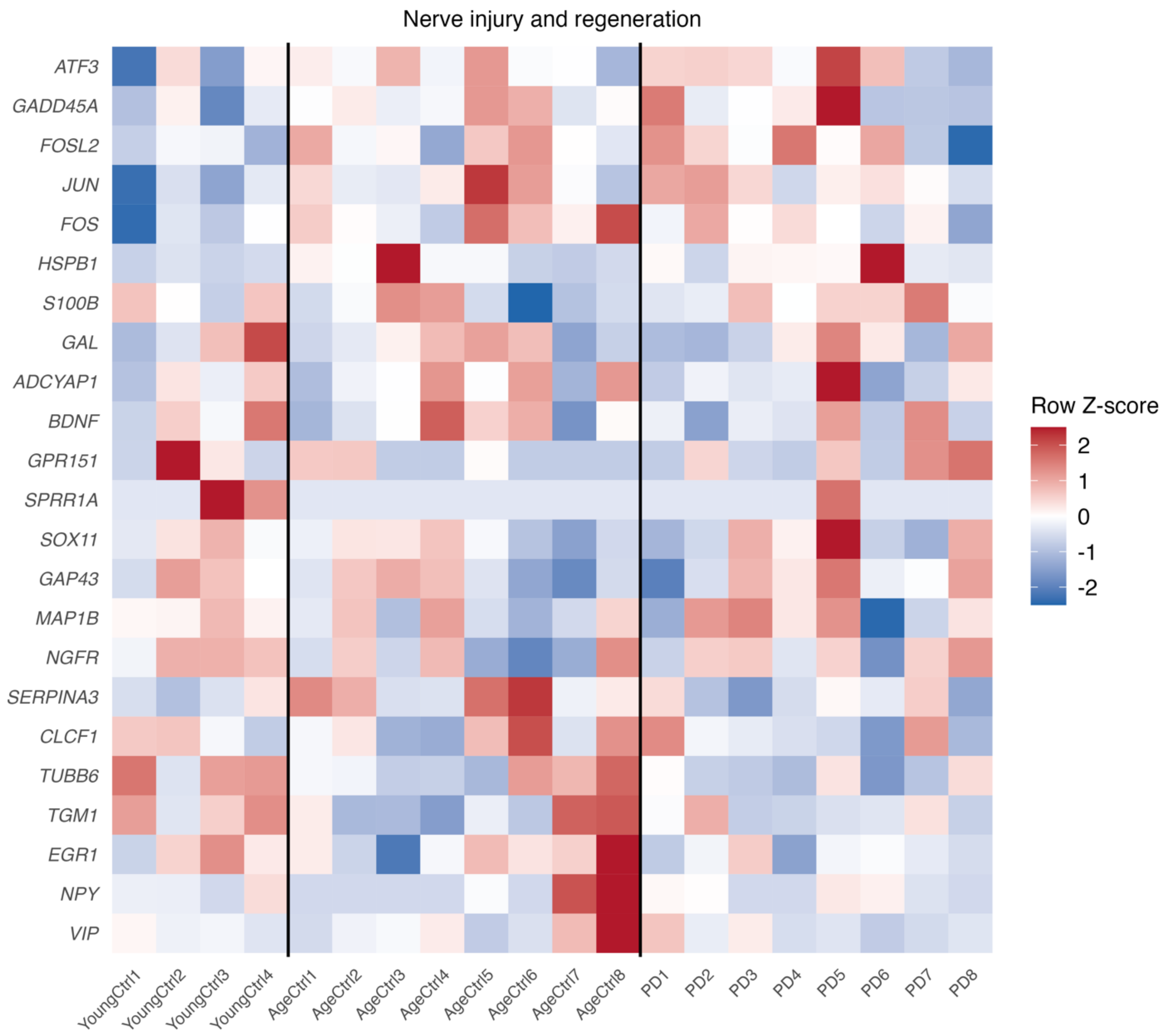


**Supplementary Figure S13. Expression patterns of nerve injury and regeneration genes across young control, age-matched control, and Parkinson's disease human dorsal root ganglia.** Heatmap showing expression of candidate genes associated with peripheral nerve injury, neuronal stress, and regenerative responses in human dorsal root ganglia (DRG) from young controls (n = 4), age-matched controls (n = 8), and individuals with Parkinson's disease (PD; n = 8). Expression values were log2(TPM + 1) transformed and standardized by row across donors to generate Z scores, with higher and lower relative expression indicated for each gene. Samples are grouped by donor cohort, with vertical lines separating young controls, age-matched controls, and PD donors. The heatmap provides a descriptive comparison of nerve injury and regeneration-associated expression patterns across age and disease groups; statistical comparisons between PD and age-matched controls were performed separately using the age-adjusted limma model.

| | Reference ligand | | | Safinamide |
|---|---|---|---|---|
| **PDB ID** | **Name** | **GlideScore (kcal/mol)** | **Redocking RMSD * (Å)** | **GlideScore (kcal/mol)** |
| 7XM9 | XEN907 | -7.543 | 1.356 | -8.451 |
| 7XMF | $Na_V$1.7-IN2 | -7.122 | 2.086 | -7.150 |
| 7XMG | TC-N1752 | -8.846 | 2.443 | -9.073 |
| 8THG | Riluzole | -6.161 | 1.428 | -7.921 |
| 8THH | Lamotrigine | -6.125 | 0.574 | -8.448 |
| * RMSD: Heavy-atom root-mean-square deviation between the top-ranked redocked pose of the reference ligand and its deposited coordinates. | | | | |

**Supplementary Table S1. Docking scores of safinamide across five $Na_V$1.7 cryo-EM structures.** For each receptor, the ligand originally resolved in that structure (reference ligand) was redocked into same binding site for benchmarking, and safinamide was independently docked into the same search space.

| Sample | Group | Age | Sex |
|---|---|---|---|
| PD1 | PD | 56 | M |
| PD2 | PD | 87 | M |
| PD3 | PD | 84 | M |
| PD4 | PD | 87 | F |
| PD5 | PD | 96 | F |
| PD6 | PD | 80 | F |
| PD7 | PD | 84 | F |
| PD8 | PD | 71 | M |
| AgeCtrl1 | Age-matched control | 90 | F |
| AgeCtrl2 | Age-matched control | 89 | M |
| AgeCtrl3 | Age-matched control | 93 | F |
| AgeCtrl4 | Age-matched control | 81 | F |
| AgeCtrl5 | Age-matched control | 80 | M |
| AgeCtrl6 | Age-matched control | 79 | M |
| AgeCtrl7 | Age-matched control | 71 | F |
| AgeCtrl8 | Age-matched control | 75 | M |
| YoungCtrl1 | Young control | 22 | F |
| YoungCtrl2 | Young control | 19 | M |
| YoungCtrl3 | Young control | 29 | F |
| YoungCtrl4 | Young control | 25 | M |

**Supplementary Table S2. Demographic characteristics of human dorsal root ganglion donors.** Demographic information for human dorsal root ganglion (DRG) samples included in the bulk RNA-sequencing analysis. Samples were obtained from individuals with Parkinson’s disease (PD; n = 8), age-matched controls (n = 8), and young controls (n = 4). Individual donor age and sex are provided for each sample.
